# Phage-delivered CRISPR-Cas9 for strain-specific depletion and genomic deletions in the gut microbiome

**DOI:** 10.1101/2020.07.09.193847

**Authors:** Kathy N. Lam, Peter Spanogiannopoulos, Paola Soto-Perez, Margaret Alexander, Matthew J. Nalley, Jordan E. Bisanz, Renuka R. Nayak, Allison M. Weakley, Feiqiao B. Yu, Peter J. Turnbaugh

## Abstract

Mechanistic insights into the role of the human microbiome in the predisposition to and treatment of disease are limited by the lack of methods to precisely add or remove microbial strains or genes from complex communities. Here, we demonstrate that engineered bacteriophage M13 can be used to deliver DNA to *Escherichia coli* within the mouse gastrointestinal (GI) tract. Delivery of a programmable exogenous CRISPR-Cas9 system enabled the strain-specific depletion of fluorescently marked isogenic strains during competitive colonization and genomic deletions that encompass the target gene in mice colonized with a single strain. Multiple mechanisms enabled *E. coli* to escape targeting, including loss of the CRISPR array or even the entire CRISPR-Cas9 system. These results provide a robust and experimentally tractable platform for microbiome editing, a foundation for the refinement of this approach to increase targeting efficiency, and a *proof-of-concept* for the extension to other phage-bacterial pairs of interest.

---

Current strategies for manipulating the microbiome either lack species- or strain-level precision^1, 2^ or require the introduction of an exogenous bacterium into the host^3, 4^. Pioneering studies of pathogenic bacteria that colonize the skin^5–7^ and gut^8^ support the potential for the use of engineered bacteriophage carrying an exogenous CRISPR-Cas system that could be directed to any target of interest; however, these methods have yet to be applied to the human or mouse microbiome. Given the tremendous diversity within both the bacterial^9^ and viral^10^ components of the human gut microbiota, we sought to establish a tripartite model system that builds upon tools for the genetic manipulation of a bacteriophage and its bacterial target coupled to an experimentally tractable mammalian host.

We focused on M13, a single-stranded DNA (ssDNA) filamentous inovirus^11, 12^ able to replicate and release virions without causing cell lysis^13^. M13 infects *E. coli* and related Enterobacteriaceae carrying the F sex factor necessary to form the conjugative F pilus^14, 15^. M13 phagemid vectors combine the advantages of plasmid DNA manipulation with the ability to easily package recombinant DNA into virions^16^. M13 has been used to target *E. coli*^17, 18^ and *Helicobacter pylori*^19^, including engineered M13 carrying CRISPR-Cas9 in an insect model of bacterial infection^5^. However, the use of M13 to deliver genetic constructs (including CRISPR-Cas systems) to cells within the mammalian gastrointestinal tract had not been previously demonstrated. We also leveraged the streptomycin (Sm)-treated mouse model for *E. coli* colonization within the mouse gut microbiota^20^, providing a robust and accessible model that could be used by any group with access to a mouse colony.

## Results

### Bacteriophage M13 enables the delivery of DNA to the gut microbiome

We utilized phagemid pBluescript II^21^ carrying the *bla* (β-lactamase) gene and a β-lactam antibiotic in the drinking water to select for successfully infected *E. coli*. pBluescript II conferred *in vitro* resistance to ampicillin and the semi-synthetic analogue carbenicillin at concentrations exceeding 1 mg/ml (**Supplementary Fig. 1**). We used streptomycin-resistant (Sm^R^) *E. coli* to colonize the GI tract of Sm-treated mice. As expected, streptomycin altered gut microbial community structure while decreasing diversity and colonization level (**Supplementary Figs. 2a-d**). Sm^R^ *E. coli* colonized at a high proportion (median 18% of the gut microbiota; range 1.4-43%) four days after gavage (**Supplementary Fig. 2e**). We introduced a Sm^R^ *E. coli* population that was a mixture of 99.9% Amp^S^ (no plasmid) and 0.1% Amp^R^ cells (pBluescript II), split the mice into two groups with access to water containing only streptomycin or both streptomycin and ampicillin, and tracked both total *E. coli* and Amp^R^ *E. coli* in mouse feces. At 6 h post-*E. coli* introduction, the percentage of Amp^R^ *E. coli* in the feces of all mice was at or close to 0.1%, consistent with the gavaged mixture transiting through the GI tract. Within 1-2 days, mice on water containing ampicillin exhibited an increase in the percent of Amp^R^ *E. coli* by 3 orders of magnitude, reaching complete or near complete colonization (**Fig. 1a**). In contrast, the Amp^R^ subpopulation was lost in mice on water without ampicillin. These results demonstrate that β-lactam antibiotics can be used to select for resistant *E. coli*.

**Figure 1.**
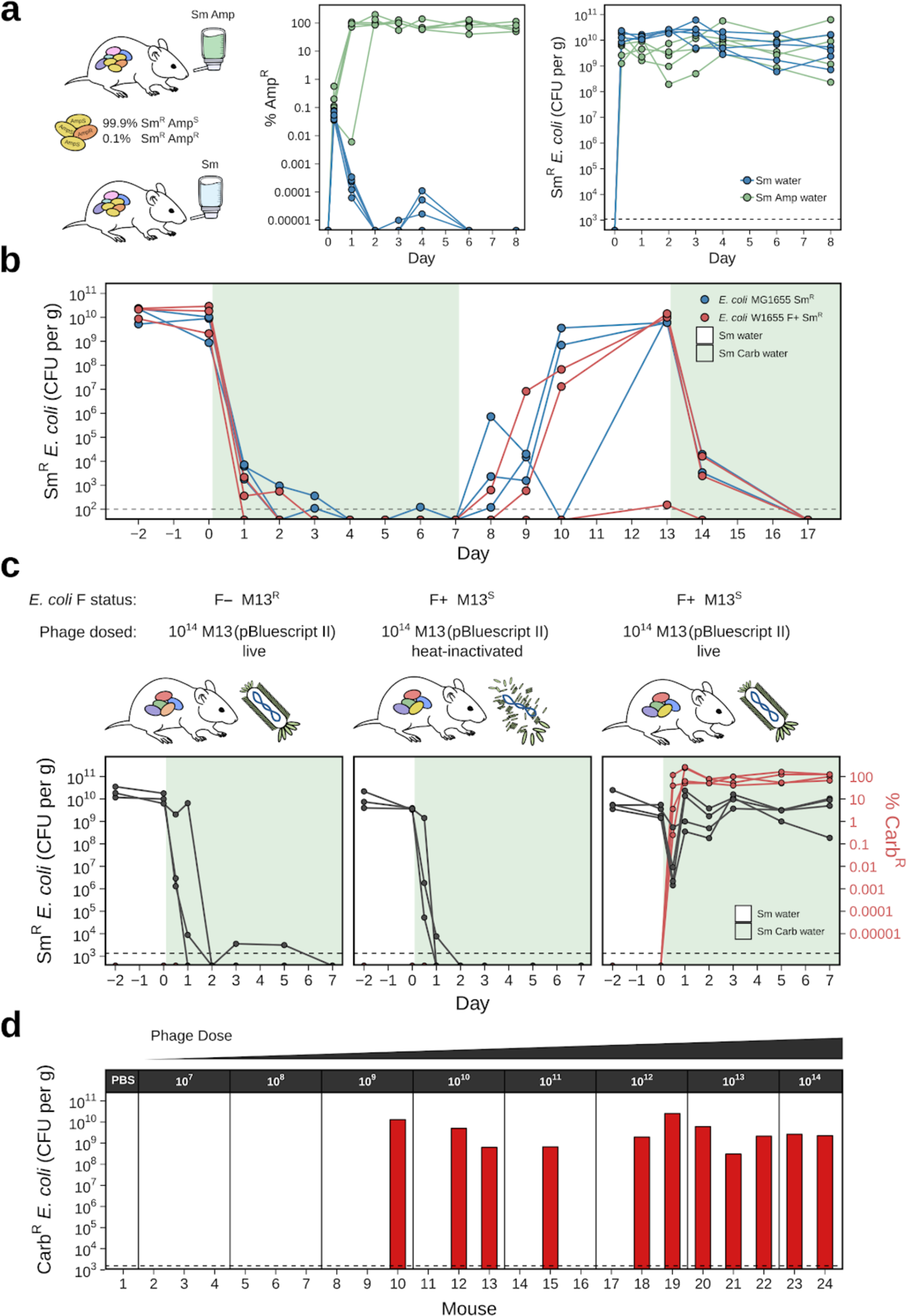
M13 bacteriophage can deliver a plasmid-borne antibiotic resistance gene to *E. coli* in the mouse gut. **(a)** A resistant subpopulation of *E. coli* can be selected in the gut when a β-lactam antibiotic is provided in the water. Streptomycin(Sm)-treated mice were gavaged with Sm^R^ MG1655 containing 99.9% ampicillin-sensitive (Amp^S^) and 0.1% ampicillin-resistant (Amp^R^) cells, and provided water containing Sm (n=5) or Sm+Amp (n=6). **(b)** A sensitive *E. coli* population is unable to maintain colonization in the gut when carbenicillin (Carb) is provided in the water. Mice were colonized with either Sm^R^ MG1655 or Sm^R^ W1655 F+ (n = 3 each) and Carb was provided in the water (shaded timepoints). **(c)** M13(pBluescript II) can infect F+ *E. coli* in the gut. Mice were split into three groups based on colonization and phage treatment: (1) Sm^R^ W1655 F− and live phage (n=3); (2) Sm^R^ W1655 F+ and heat-inactivated phage (n=3); (3) Sm^R^ W1655 F+ and live phage (n=4). 10^14^ phage were dosed and Carb was provided in the water. Total *E. coli* (black) and %Carb resistant colonies (Carb^R^; red) in mouse fecal pellets are shown. **(d)** M13-based delivery of an antibiotic resistance gene is dose-dependent. Varying doses (10^7^-10^14^) of M13(pBluescript II) were given to mice (n=2-3/dose) and Carb was provided in the water. After 2 days, Carb^R^ CFU/gram feces was determined. Dashed line indicates our limit of detection.

**Figure 2.**
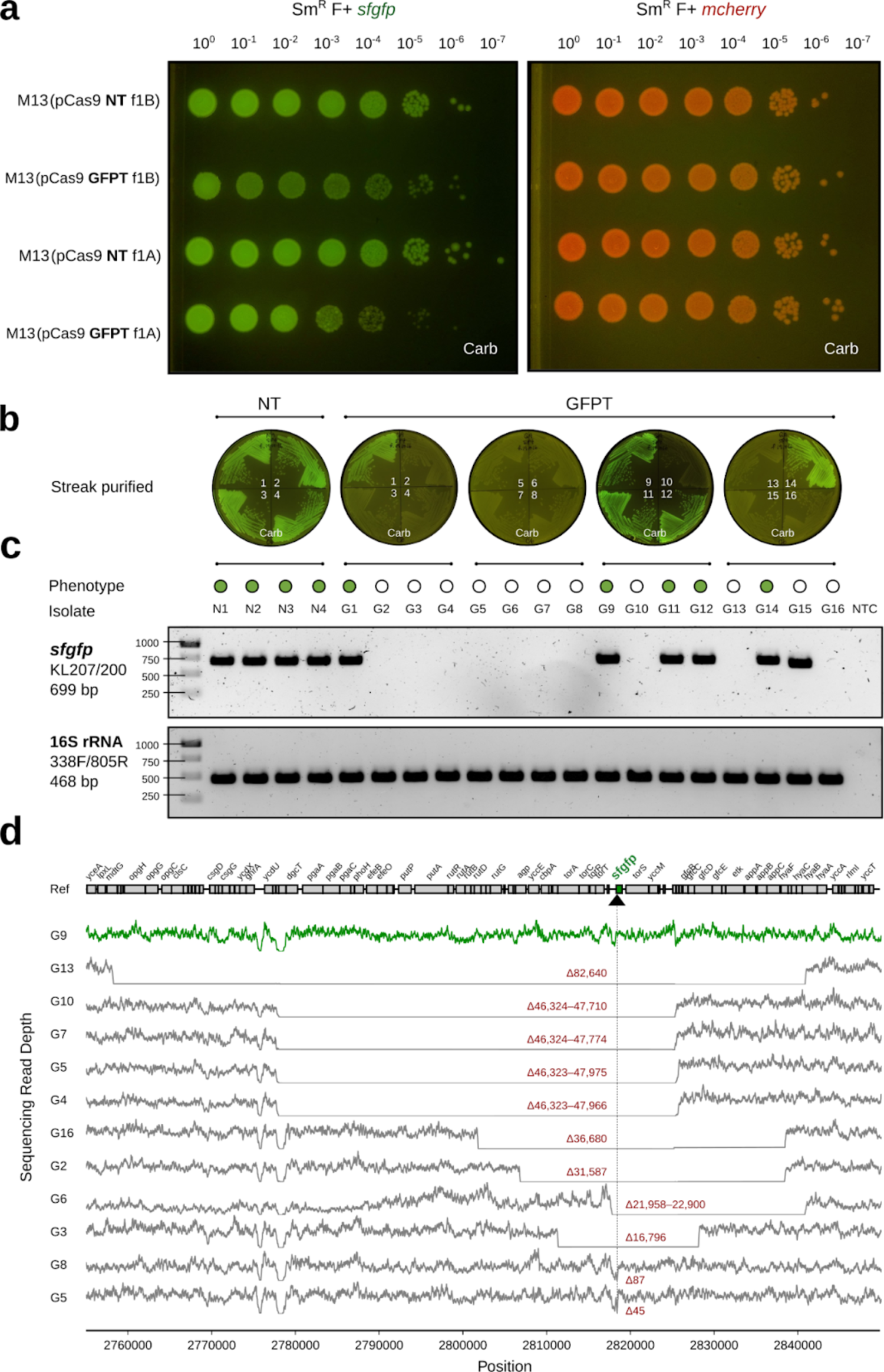
M13-mediated delivery of CRISPR-Cas9 to *E. coli in vitro* causes impaired colony growth and can induce chromosomal deletions that encompass the targeted gene. **(a)** GFP+ *E. coli* exhibit a sick colony morphology after infection with M13 phage carrying GFP-targeting CRISPR-Cas9. NT (non-targeting) or GFPT (GFP-targeting) M13 were used to infect Sm^R^ W1655 F+ *sfgfp* or Sm^R^ W1655 F+ *mcherry* as a control. Cells were infected, diluted, and spotted onto media with selection for the vector; f1A or f1B indicates version of vector. **(b)** CRISPR-Cas9 targeting the *sfgfp* gene can induce loss of fluorescence. Colonies arising from infection with NT-M13 or GFPT-M13 were subjected to several rounds of streak purification on selective media to ensure phenotypic homogeneity and clonality. The majority (11/16) of GFPT clones exhibited a loss of fluorescence. **(c)** Clones exhibiting loss of fluorescence either lack an *sfgfp* PCR amplicon or exhibit an amplicon of decreased size. Genomic DNA was isolated from streak-purified clones and PCR was used to determine whether the *sfgfp* gene was present; PCR for the 16S rRNA gene was performed as a positive control. **(d)** Genome sequencing results confirm that nonfluorescent clones have chromosomal deletions encompassing the targeted gene. Read depth surrounding *sfgfp* locus for G9 clone (green line, fluorescent control) and all nonfluorescent clones (grey lines). Deletion size indicated in red; range indicates a deletion flanked by repetitive sequences. Black arrow and vertical line denote position of targeting. Carb, carbenicillin.

Antibiotics were capable of eradicating a sensitive population of *E. coli* that had established stable colonization in the mouse gut. We colonized Sm-treated mice with Sm^R^ *E. coli* MG1655 or W1655 F+ and tracked colonization levels during treatment with the β-lactam antibiotic carbenicillin. Carbenicillin decreased the median *E. coli* colonization level from 9.6 × 10^9^ to 2.0 × 10^3^ CFU/gram feces in the first day, and levels decreased to below our limit of detection (∼10^2^ CFU/g) in all mice over the course of treatment (**Fig. 1b**). When selection was lifted on Day 7, recolonization was observed for 5/6 mice. When carbenicillin was reintroduced on Day 13, colonization again dropped below our limit of detection. The low background of *E. coli* in the gut during carbenicillin treatment, as well as the lack of spontaneous resistant cells able to expand, supports the utility of this model for assessing the phage-mediated delivery of a resistance gene.

Next, we tested our ability to deliver an antibiotic resistance gene to *E. coli* within the gut. We colonized Sm-treated mice with either Sm^R^ *E. coli* W1655 F+ (M13^S^) or W1655 F− (M13^R^ as a control), and dosed each animal with either live or heat-inactivated M13 carrying pBluescript II (**Fig. 1c**). After dosing the mice with 1×10^14^ M13(pBluescript II), we immediately transferred them to water containing carbenicillin and tracked both total *E. coli* and Carb^R^ *E. coli* in the feces. *E. coli* colonization fell rapidly and stayed near or below the limit of detection in control mice that were either colonized with F− and given live phage or colonized with F+ but given heat-inactivated phage. In contrast, mice colonized with F+ and dosed with live phage had a transient drop in colonization on the first day, during which the rise of Carb^R^ cells occurred, and colonization was re-established within one day by an *E. coli* population resistant to carbenicillin (**Fig. 1c**). These results suggest that orally dosed M13 phage were able to infect *E. coli* in the gut and deliver a plasmid conferring resistance to carbenicillin.

We replicated M13-mediated pBluescript II delivery to *E. coli* in the gut in an independent animal experiment. Sm-treated mice were colonized with Sm^R^ *E. coli* W1655 F+ and orally dosed with ten-fold serial dilutions of M13(pBluescript II). Colonization by Carb^R^ *E. coli* was consistent at high doses but variable at lower doses representing a significantly higher probability of successful colonization with increasing phage dose (**Fig. 1d**; P=0.009, odds ratio=2.5, logistic regression). Plasmid DNA of the expected size was detected in fecal Carb^R^ *E. coli* isolates from all 11 mice that were successfully colonized (**Supplementary Fig. 3a**). Genome sequencing confirmed the presence of pBluescript II in these 11 isolates, which was undetectable in the parent strain (**Supplementary Fig. 3b**). These results indicate that plasmid DNA was transferred from M13 phage into recipient *E. coli* colonizing the GI tract.

Finally, we repeated this experiment in the absence of carbenicillin selection. We colonized mice with Sm^R^ *E. coli*, gavaged each mouse with M13(pBluescript II), and tracked both infected (Carb^R^) and total (Sm^R^) *E. coli* in feces. The fraction of phage-infected Carb^R^ *E. coli* was low, reaching a maximum of 0.1% of the total population (**Supplementary Fig. 4**), potentially indicative of poor phage survival during GI transit. We gavaged mice with M13(pBluescript II) and assayed for viable phage in the feces. The median output of viable M13(pBluescript II) was reduced to 1×10^6^ relative to an input of 6×10^13^ (**Supplementary Figs. 5a,b**). M13(pBluescript II) is acid tolerant *in vitro* (**Supplementary Fig. 5c**), suggesting that additional factors may be responsible for the low *in vivo* viability and emphasizing the benefits of pairing gene delivery with antibiotic selection.

### M13 carrying CRISPR-Cas9 can target *E. coli in vitro*

We generated two fluorescently marked isogenic derivatives of Sm^R^ W1655 F+ using the *mcherry* (red fluorescence) or the *sfgfp* (green fluorescence) marker gene. Next, we constructed M13-compatible non-targeting (NT) and GFP-targeting (GFPT) CRISPR-Cas9 vectors by cloning the spacers sequences, *bla* gene, and f1 origin of replication into the previously described low-copy vector pCas9^22^, generating pCas9-NT-f1A/B and pCas9-GFPT-f1A/B (**Supplementary Fig. 6**). The *bla* and f1 *ori* were cloned as a fragment from pBluescript II in both possible orientations (A or B) to make possible M13 ssDNA packaging of either strand of vector DNA. We packaged these phagemids into M13 using a helper strain and called the resulting phage NT-M13 or GFPT-M13. The two phage were used to infect the GFP+ or mCherry+ strains and cells were diluted and spotted on solid media containing carbenicillin to select for the transferred phagemid. GFP+ *E. coli* infected with GFPT-M13 exhibited impaired colony growth relative to the NT-M13 control (**Fig. 2a** and **Supplementary Fig. 7**). Total CFUs were not markedly affected, indicating that cells can recover from M13-delivered CRISPR-Cas9 targeting.

Analysis of the surviving cells provided mechanistic insights. Colonies arising from infection with NT-M13 or GFPT-M13 were streak purified, allowing us to pick a mixture of bright and dim colonies. Of 16 GFPT clones analyzed, 11 were non-fluorescent (**Fig. 2b**). PCR amplification of *sfgfp* confirmed the intact gene in 4 NT controls and all 5 GFPT clones that retained fluorescence (**Fig. 2c**). Sanger sequencing revealed that 1 GFPT clone had a point mutation in the *sfgfp* target (**Supplementary Fig. 8**) while the 4 others had lost the spacer in the CRISPR-Cas9 phagemid that leads to targeting (**Supplementary Fig. 9**). All 11 non-fluorescent GFPT clones retained the spacer (**Supplementary Fig. 9**) and had chromosomal deletions at the target locus: 10 were PCR negative (**Fig. 2c**) and 1 had a small deletion within *sfgfp* (**Supplementary Fig. 8**). Finally, we used whole genome sequencing to define the size of each deletion, which ranged from 45 bp to 82.6 kb (**Fig. 2d**), consistent with prior work demonstrating that *E. coli* can repair Cas9-induced double-stranded breaks through homologous recombination^23^.

These results led us to hypothesize that targeted cells would be less able to recover during competitive growth. We co-cultured GFP+ and mCherry+ *E. coli*, adding either NT-M13 or GFPT-M13 followed by carbenicillin to select for phage infection. GFPT-M13 decreased the frequency of GFP+ colonies by 4 h, relative to the NT-M13 control (**Fig. 3a**). At later timepoints (16-24 h), healthy GFP+ colonies increased in abundance, consistent with low levels of carbenicillin after 4 h in cultures expressing the β-lactamase resistance gene (**Fig. 3b**). We confirmed the loss of selection for the phagemid by re-analyzing our colonies on selective media (**Supplementary Fig. 10**). Next, we used flow cytometry to better quantify the two strains in an independent experiment. Compared to the NT-M13 control, GFPT-M13 co-cultures exhibited fewer GFP+ events (**Fig. 3c**) and a bimodal distribution of fluorescence (**Fig. 3d**). Counts of GFP+ cells were higher by flow cytometry than on solid media for the same co-cultures (**Fig. 3c** inset), consistent with an impaired growth of these cells. GFP+ events further decreased at 24 h in the GFPT-M13 group (**Supplementary Fig. 11**). Taken together, these results suggest that competitive growth can increase the efficiency of targeting a strain for depletion due to the resulting growth impairments in the targeted strain.

**Figure 3.**
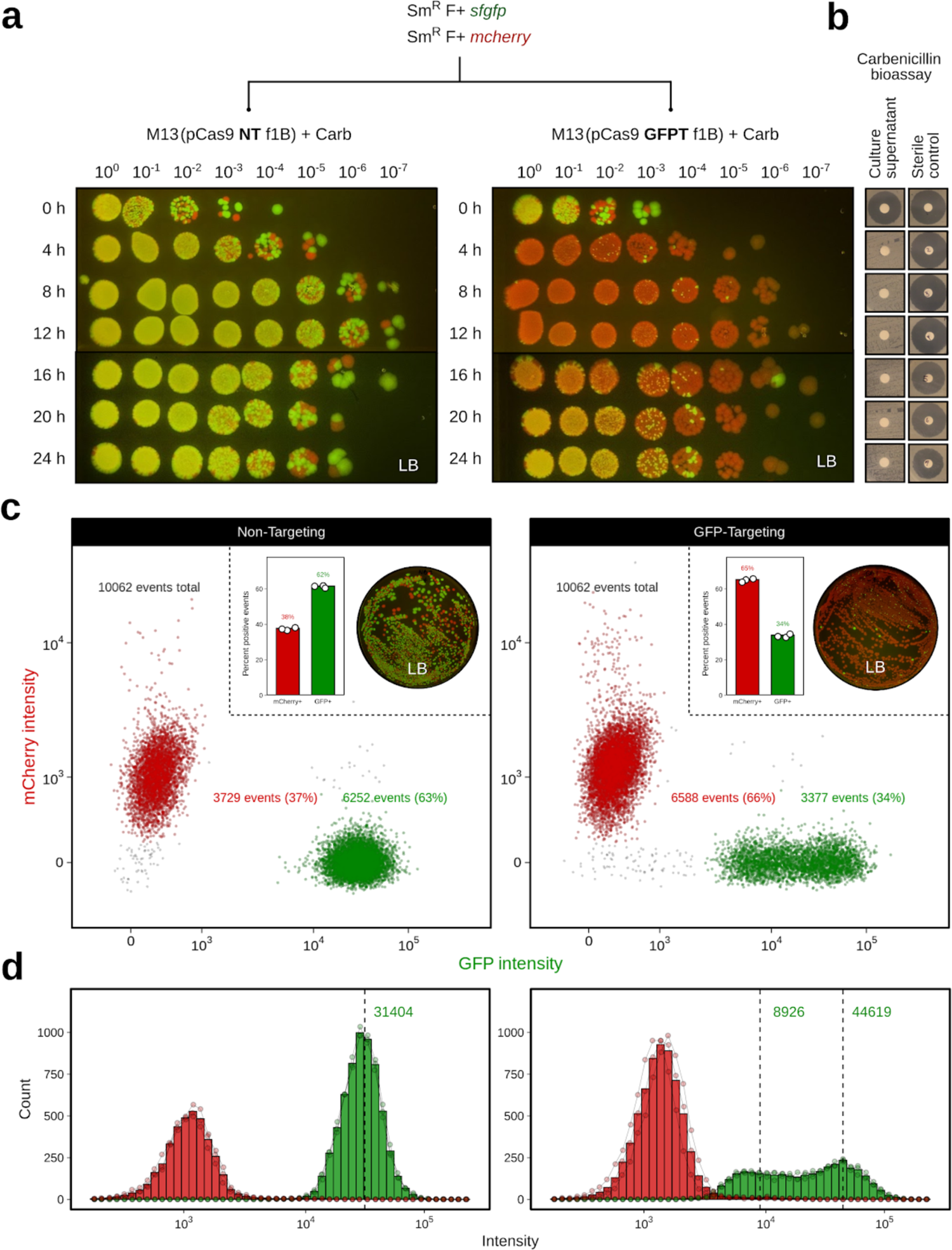
M13-delivered CRISPR-Cas9 for sequence-specific targeting of *E. coli* in *in vitro* co-cultures of fluorescently marked isogenic strains. **(a)** M13-delivered GFP-targeting CRISPR-Cas9 leads to reduced competitive fitness of the GFP-marked strain. A co-culture of Sm^R^ F+ *sfgfp* and Sm^R^ F+ *mcherry* was incubated with NT-M13 or GFPT-M13 at a starting MOI of ∼500. Carbenicillin (Carb) was added to a final concentration of 100 µg/ml to select for phage infection. Co-cultures were sampled every 4 h over 24 h; cells were washed, serially diluted, and spotted onto non-selective media to assess targeting of the GFP-marked strain. Carbenicillin in culture supernatants was not detectable within 4 h of growth, using a carbenicillin bioassay against indicator strain *Bacillus subtilis* 168; bioassay detection limit approximately 2.5 µg/ml. **(c)** Flow cytometry of co-cultures 8 h following the addition of phage and carbenicillin show reduced GFP+ events in the GFPT versus NT condition. Representative flow plot shows data from one of three replicates. Inset: bar graph quantifying percent GFP+ and mCherry+ events for three replicates (left); plating results for a single replicate on non-selective media (right). **(d)** GFPT CRISPR-Cas9 changes the shape of the distribution of GFP+ population. Histogram of mCherry+ and GFP+ events by intensity shows that a proportion of GFP+ cells in the GFPT condition have shifted to a state of lower fluorescence. Bars indicate the mean of three replicates; connected points are individual replicates.

### Sequence-specific depletion of *E. coli* within the mouse gut microbiota

We co-colonized Sm-treated mice with both Sm^R^ F+ *sfgfp* and Sm^R^ F+ *mcherry* strains, orally dosed them with either 10^11^ NT-M13 or GFPT-M13, and added carbenicillin in the water to select for phage infection. After one week of treatment, carbenicillin was removed from the water and mice were followed for an additional week to determine whether phage-induced changes would persist in the absence of maintaining selection (**Fig. 4a**). Flow cytometry on mouse stool samples revealed that the GFP+ strain outcompeted the mCherry+ strain in the NT-M13 group (**Figs. 4b,c** and **Supplementary Figs. 12,13**). In contrast, GFP+ events in the GFPT-M13 group exhibited a sharp decrease on Day 2, followed by a recovery on Days 7 and 14 to levels below the NT-M13 group (**Figs. 4b,c**). Culturing from mouse stool confirmed the decreased GFP+ events on Day 2 (**Supplementary Fig. 14**). In 4 mice that received GFPT-M13, the mCherry+ strain fixed in the population (GFP+ events were below background), an outcome that was not observed for any mouse in the NT-M13 group (**Fig. 4d**). Despite lifting the carbenicillin selection for 1 week, endpoint GFP+ events remained significantly lower in the GFPT-M13 group relative to NT-M13 controls (**Fig. 4e**; P=0.0002, Mann-Whitney test). These data support the utility of M13-delivered CRISPR-Cas9 for sequence-specific depletion of an otherwise isogenic bacterial strain in the mouse gut.

**Figure 4.**
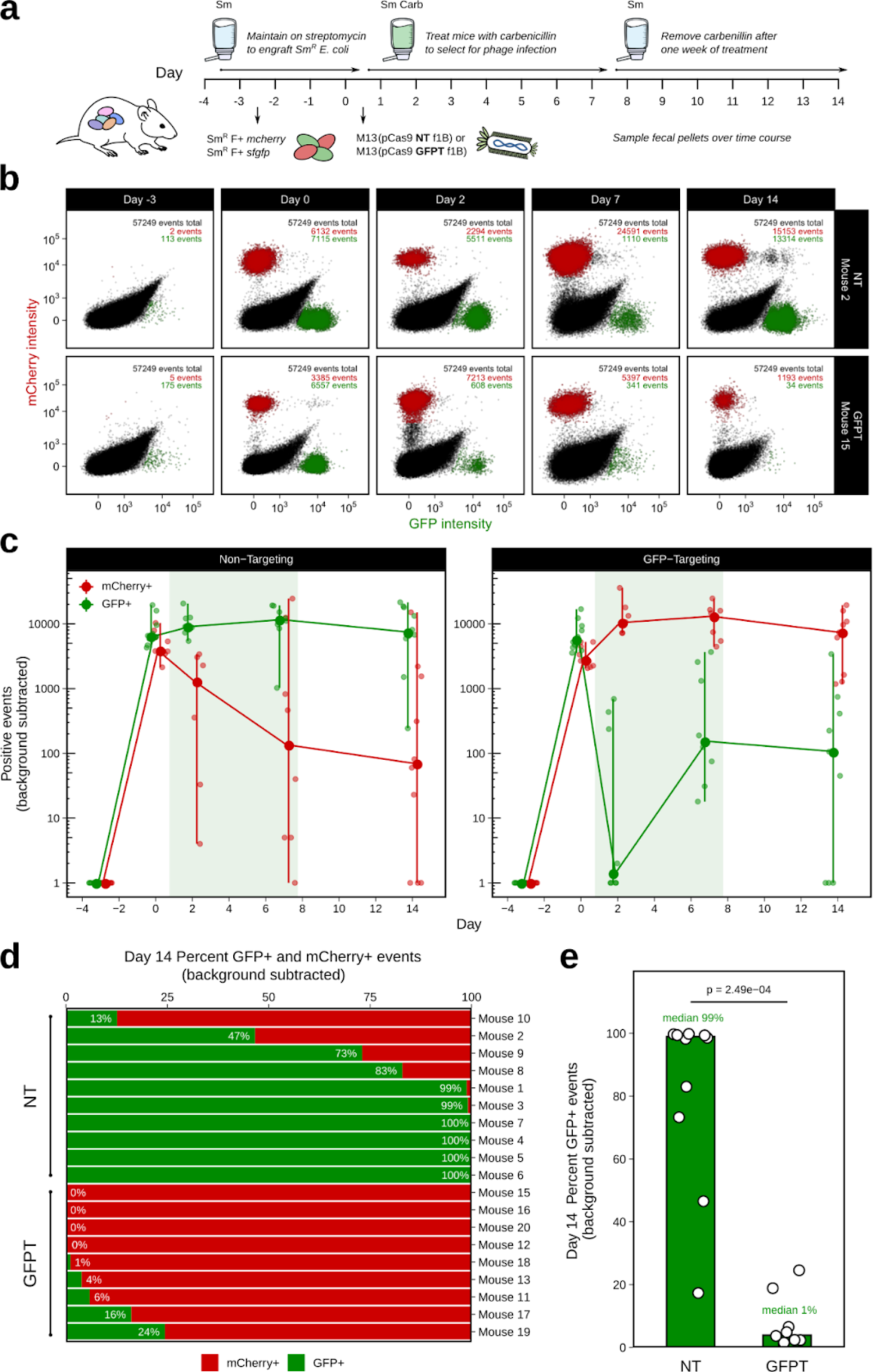
M13-delivered CRISPR-Cas9 for sequence-specific depletion of *E. coli* in the gut of mice colonized by competing fluorescently marked isogenic strains. (a) **Timeline: Day** −3, colonize with 50/50 mixture of streptomycin(Sm)-resistant Sm^R^ F+ *sfgfp* and Sm^R^ F+ *mcherry*; Day 0, dose with 10^11^ NT-M13 or GFPT-M13 (n=10/group) and provide carbenicillin (Carb) in the water; Day 7, remove carbenicillin. **(b)** GFPT-M13 can lead to loss of the GFP-marked strain. Time series flow plots of fecal samples for one mouse from each of NT and GFPT groups. Top right: number of total, mCherry+, and GFP+ events. **(c)** Mice in GFPT group exhibited a decrease in number of fecal GFP+ events in over time compared to NT group; timepoints were excluded if both GFP+ and mCherry+ events were below background thresholds. Line graph: points indicate median; vertical lines, range. **(d)** Mice in GFPT group exhibited depletion or loss of the GFP-marked strain. Percent GFP+ and mCherry+ events for each mouse on Day 14. Mice were excluded if both GFP+ and mCherry+ events were both below background thresholds (final n=9 GFPT and n=10 NT). **(e)** A significant difference was observed in the percent of GFP+ events in fecal samples at Day 14 in the GFPT group compared to NT. Bars are medians; *p*-value, Mann-Whitney test.

### M13-delivered CRISPR-Cas9 induces chromosomal deletions in the gut microbiome

We constructed a double-marked Sm^R^ F+ *sfgfp mcherry* strain to quantify the efficiency of gene deletion. We introduced this strain into Sm-treated mice, orally dosed each mouse with either 10^11^ NT-M13 or GFPT-M13, and added carbenicillin in the water. After one week, we removed carbenicillin and followed the mice for a final week (**Fig. 5a**). GFP-mCherry+ events were detectable in GFPT-M13 but not NT-M13 mice, indicative of successful CRISPR-Cas9 delivery and gene deletion (**Fig. 5b** and **Supplementary Fig. 15**). By the final timepoint, GFP-mCherry+ events were detected in 3/8 mice (**Fig. 5c** and **Supplementary Fig. 16a**). The relative abundance of GFP-mCherry+ cells varied from 12-96% (**Fig. 5c**). Culturing on solid media confirmed the presence of viable red fluorescent colonies in proportions consistent with flow cytometry results (**Fig. 5d** and **Supplementary Fig. 16b**).

**Figure 5.**
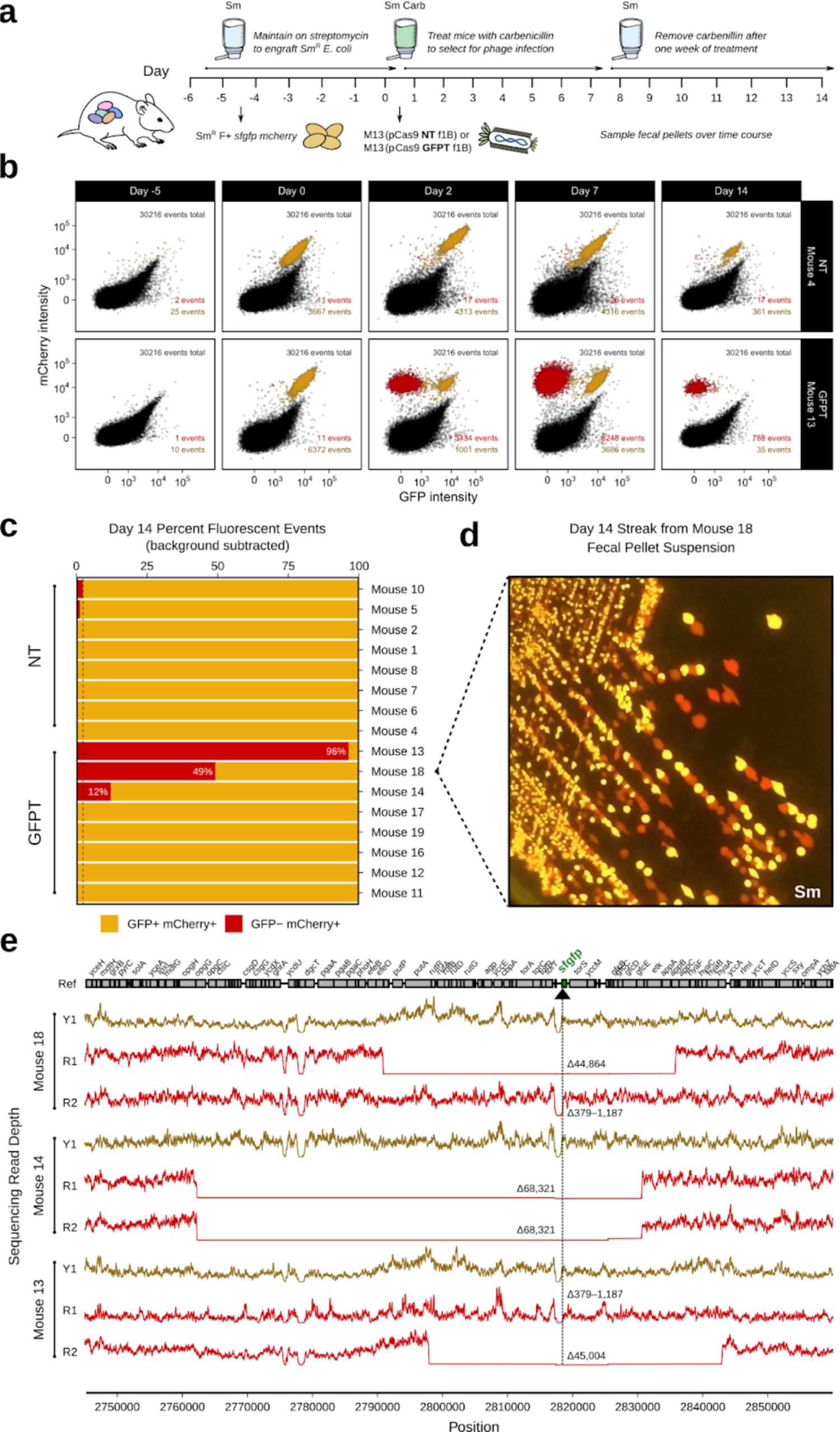
M13-delivered CRISPR-Cas9 can induce chromosomal deletions encompassing the targeted gene in *E. coli* colonizing the mouse gut. **(a)** Timeline: Day-5, colonize with double-marked streptomycin(Sm)-resistant Sm^R^ F+ *sfgfp mcherry*; Day 0, dose with 10^11^ NT-M13 or GFPT-M13 (n=10/group) and provide carbenicillin (Carb) in the water; Day 7, remove carbenicillin. **(b)** GFPT-M13 can cause loss of GFP fluorescence in double-marked *E. coli*. Time series flow plots of fecal samples for select mice, one from each of NT and GFPT groups. Top right: total number of events; bottom right: GFP-mCherry+ events and GFP+ mCherry+ events. Day 14 fecal samples of three mice in GFPT group exhibited mCherry-only fluorescence. Percent GFP+ mCherry+ and GFP-mCherry+ events for each mouse; mice were excluded if both populations were below the background threshold (final n=8/group). Dashed line indicates maximum mCherry fluorescence for NT group. **(d)** Colonies arising from culture of Mouse 18 Day 14 fecal sample confirmed presence of red-only fluorescence. **(e)** Genome sequencing results confirm red fluorescent isolates from Mouse 13, 14, and 18 have chromosomal deletions encompassing the targeted gene. Read depth surrounding *sfgfp* locus for GFP+ mCherry+ (Y1; yellow lines) and GFP-mCherry+ fluorescent (R1 or R2; red lines) isolates from Day 2 fecal samples. Deletion size indicated in red; range indicates a deletion flanked by repetitive sequences. Black arrow and vertical line denote position of targeting.

To more definitively assess the presence or absence of the targeted genomic locus and the CRISPR-Cas system, we isolated GFP-mCherry+ and GFP+ mCherry+ *E. coli* from Day 2 mouse stool. All of the GFP+ mCherry+ isolates from the NT-M13 group and the GFP-mCherry+ isolates from the GFPT-M13 group had an intact spacer sequence (**Supplementary Fig. 17a**). In contrast, 4/5 GFP+ mCherry+ isolates from the GFPT-M13 group had lost the spacer (**Supplementary Fig. 17a**). Of note, the remaining isolate lost *cas9*, and parts of the CRISPR array and tracrRNA (**Supplementary Figs. 17b,c**). Whole genome sequencing was used to confirm putative chromosomal deletions and to quantify their size. Two representative colonies were analyzed from each of the 3 mice with detectable GFP-mCherry+ cells, revealing a wide range in deletion sizes that were not observed in a control GFP+ mCherry+ isolate from each animal (**Fig. 5e**). These results indicate that while it is possible for CRISPR-Cas9-induced genomic deletion events to occur *in vivo*, resultant deletion strains may or may not outcompete the parent strain due to the potential to evade targeting through loss of some or all of the exogenous CRISPR-Cas system.

## Discussion

Our results emphasize that foundational, reductionist, and highly controlled studies will be necessary to assess the feasibility, utility, and limitations of phage-based gene delivery as a tool for microbiome editing. While our results provide a *proof-of-principle* for strain-specific targeting within the GI tract, the full eradication of the targeted strain was difficult to achieve due to the ability of bacterial cells to survive Cas9-induced double-stranded breaks by homologous recombination^24^. We propose that CRISPR-Cas9 may be better suited to induce targeted genomic deletions, leveraging the conserved DNA repair pathways present in bacteria. An advantage of this approach is that the deletion of a single genomic locus is unlikely to have as large an impact on the rest of the gut microbiota than if the strain were to be removed entirely. Remarkably, we detected a wide range of deletion sizes (379–68,321 bp), highlighting the ability of bacteria to survive large deletions and opening up the potential for the *in vivo* removal of entire biosynthetic gene clusters or pathogenicity islands. In turn, our data suggests that it may also be feasible to deliver more complex genetic circuits to *E. coli*, with the goal of boosting metabolic pathways beneficial to its mammalian host. Furthermore, the size of resulting chromosomal deletions could be controlled by providing a DNA repair template alongside CRISPR-Cas9^22^.

There are several limitations of our current approach that could be refined through iterative “design-build-test” cycles. Perhaps most importantly, we identified multiple mechanisms through which cells can escape CRISPR-Cas9 targeting during *in vitro* growth and/or within the GI tract, including the loss of the spacer in the CRISPR array, target site mutations, and even one case in which the entire CRISPR-Cas9 system was deleted. Spacer loss could be reduced through the use of single guide RNAs (sgRNAs)^25^ or through the mutation of the repeat sequences that flank each spacer^26^. Target site mutation could be countered by targeting multiple sites simultaneously and by prioritizing conserved regions of the target genes essential for activity.

Another important caveat is that antibiotics were used to select for successful gene delivery, which is not ideal due to potential disruptions to the gut microbiota^1^ and/or selection for antibiotic resistance. Removing this selection resulted in a low penetrance of cells carrying the delivered cargo, similar to that observed for delivery to bacteria in a soil community in the absence of selection^27^. Low rates of gene delivery are likely driven by the observed loss of viable M13 bacteriophage during transit through the GI tract. We confirmed that M13 is acid tolerant; however, it is sensitive to artificial gastric juice^28^, suggesting that gastric proteases may be responsible for low phage survival. The aggregative behaviour of filamentous phage in response to microbial and/or host polymers^29^ may also be important to consider. Encapsulating phage for oral transit may be able to circumvent these limitations or alternatively, it may be possible to use non-antibiotic selection strategies such as dietary supplementation of an exclusive substrate^3^ that can only be used by phage-infected strains.

In conclusion, this work provides a valuable step towards establishing a modular toolkit for microbiome editing. The extension of these approaches to enable the genetic manipulation of a more diverse panel of bacteria found within the human microbiota will require a renewed effort to isolate and characterize bacteriophage that target strains of interest^30^. Robust *in vitro* methods to study and genetically modify novel bacteriophage are also needed, given that most of their host bacteria remain genetically intractable^31^. Finally, our work emphasizes the value of model systems for understanding the rules of engagement, complementing ongoing efforts to advance candidate phage-based therapies into the clinic.

## Methods

### Strains, plasmids, phage, and oligonucleotides

Bacterial strains, plasmids, and phage used in this study, including descriptions and sources, are provided in **Supplementary Table 1**. Oligonucleotides used in this study are provided in **Supplementary Table 2**.

### Minimum inhibitory concentration (MIC) assay

Cells were prepared by standardizing an overnight culture to an OD_600_ of 0.1 using saline (0.85% NaCl), and further diluted ten-fold in saline then ten-fold in LB. The drug was prepared by dissolving the antibiotic in vehicle (sterile distilled water) and filter-sterilizing, then serially diluting two-fold in vehicle to prepare 100× stock solutions, and finally diluting ten-fold in LB for 10× stock. To wells of a 96-well plate, 60 µl of LB, 15 µl of drug, and 75 µl of cells were added and mixed well. Final drug concentrations ranged between 0.002 µg/ml to 1000 µg/ml for ampicillin and 0.24 µg/ml to 2000 µg/ml for carbenicillin. The plate was incubated overnight at 37°C without shaking and OD_600_ was measured the following morning after agitation.

### 16S rRNA gene sequencing

Mouse fecal pellets were stored at −80°C. DNA was extracted from single pellets using a ZymoBIOMICS 96 MagBead DNA Kit and 16S rRNA gene sequencing was performed using a dual indexing strategy^32^. Briefly, a 22-cycle primary PCR was performed using KAPA HiFi Hot Start DNA polymerase (KAPA KK2502) and V4 515F/806R Nextera primers. The reaction was diluted in UltraPure DNase/RNase-free water (Life Tech 0977-023) and used as template for a 10-cycle secondary (indexing) PCR using sample-specific dual indexing primers. The reactions were normalized using a SequelPrep Normalization plate (Life Tech A10510-01) and the DNA was eluted and pooled. To purify and concentrate the DNA, 5 volumes of PB Buffer (Qiagen 28004) were added, mixed, and purified using a QIAquick PCR Purification Kit (Qiagen 28106). The DNA was gel extracted using a MinElute Gel Extraction Kit (Qiagen 28604), quantified by qPCR using a KAPA Library Quantification Kit for Illumina Platforms (KAPA KK4824), and paired-end sequenced on the Illumina MiSeq platform. Data were processed using a 16S rRNA gene analysis pipeline (https://github.com/jbisanz/AmpliconSeq) based on QIIME2^33^ incorporating DADA2^34^, and analyzed using R packages qiime2R (v0.99.23; https://github.com/jbisanz/qiime2R), phyloseq (v1.33.0)^35^, and phylosmith (v1.0.4)^36^. See Data Availability for more information.

### Construction of streptomycin-resistant *E. coli* strains

Strains resistant to the antibiotic streptomycin were generated by either selection for spontaneous resistance or by lambda Red recombineering^37, 38^. Spontaneous resistant mutants were selected by plating overnight cultures on LB supplemented with 500 µg/ml streptomycin. Lambda Red recombineering was later used to introduce a specific allele for genetic consistency between strains as different mutations in the *rpsL* gene can confer resistance to streptomycin^39^. Briefly, cells were transformed with the Carb^R^ temperature-sensitive plasmid pSIJ8^38^, and electrocompetent cells were prepared from cells grown in LB carbenicillin at 30°C to early exponential phase and lambda Red recombinase genes were induced by addition of L-arabinose to 7.5 mM. Cells were electroporated with an *rpsL*-Sm^R^ PCR product amplified from a spontaneous streptomycin-resistant mutant of MG1655 using primers PS-rpsL1 and PS-rpsL2, and recombinants were selected on LB supplemented with 500 µg/ml streptomycin. The pSIJ8 plasmid was cured by culturing in liquid at 37°C in the absence of carbenicillin, plating for single colonies, and confirming Carb^S^. The *rpsL* gene of Sm^R^ strains was confirmed by Sanger sequencing.

### Construction of fluorescently marked *E. coli* strains

P1 lysates were generated of AV01::pAV01 and AV01::pAV02 carrying clonetegrated *sfgfp* and *mcherry*, respectively^40^. Briefly, 150 µl of overnight culture in LB supplemented with 12.5 µg/ml kanamycin was mixed with 1 µl to 25 µl P1 phage (initially propagated from ATCC on MG1655). The mixture was incubated for 10 min at 30°C to aid adsorption, added to 4 ml LB 0.7% agar, and overlaid on pre-warmed LB agar supplemented with 25 µg/ml kanamycin 10 mM MgSO_4_. Plates were incubated overnight at 30°C, and phage were harvested by adding 5 ml SM buffer, incubating at room temperature for 10 min, and breaking and scraping off the top agar into a conical tube. Phage suspensions were centrifuged to pellet agar; the supernatant was passed through a 100 µm cell strainer, then through a 0.45 µm syringe filter, and lysates were stored at 4°C. For transduction, 1-2 ml of recipient overnight culture was pelleted and resuspended in 1/3 volume LB 10 mM MgSO_4_ 5 mM CaCl_2_. 100 µl of cells was mixed with 1 µl to 10 µl P1 lysate and incubated at 30°C for 60 minutes. To minimize secondary infections, 200 µl 1 M sodium citrate was added, followed by 1 ml of LB. The mixture was incubated at 30°C for 2 h, then plated on LB 10 mM sodium citrate 25 µg/ml kanamycin to select for transductants. For excision of the vector backbone including the kanamycin resistance gene and heat-inducible integrase, cells were electroporated with pE-FLP^41^; transformants were selected on carbenicillin and confirmed for Km^S^. pE-FLP was cured by culturing in liquid at 37°C in the absence of carbenicillin, plating for single colonies, and confirming Carb^S^. Strains were subsequently grown routinely at 37°C. For imaging fluorescent strains on agar, plates were typically incubated at 37°C overnight, transferred to room temperature to allow fluorescence intensity to increase, and then imaged.

### Mouse experiments with *E. coli* colonization, antibiotic water, and phage treatment

Animal procedures were approved by the University of California, San Francisco (UCSF) Institutional Animal Care and Use Committee (IACUC), and animal experiments performed were in compliance with ethical regulations. Specific pathogen free female BALB/c mice from the vendor Taconic were used for all mouse experiments. Streptomycin water was prepared by dissolving USP grade streptomycin sulfate (VWR 0382) in autoclaved tap water to a final concentration of 5 mg/ml and passing through 0.45 µm filtration units. Mice were provided streptomycin water for 1 day, followed by oral gavage of 0.2 ml containing approximately 10^9^ CFU of streptomycin-resistant *E. coli*. Mice were kept on streptomycin water thereafter to maintain colonization. For selection with β-lactam antibiotics, USP grade ampicillin sodium salt (Teknova A9510) or USP grade carbenicillin disodium salt (Teknova C2110) was also dissolved in the water to a final concentration of 1 mg/ml; carbenicillin was preferred for its increased stability over ampicillin^42^. Drinking water containing streptomycin was prepared fresh weekly; with the addition of a β-lactam antibiotic, it was prepared fresh every 3-4 days. For phage treatment, filtered phage solutions stored at −80°C were thawed and used directly for oral gavage. Unfiltered phage solutions were precipitated by diluting approximately 5-fold in PBS, adding 0.2 volumes phage precipitation solution (20% PEG-8000, 2.5 M NaCl), incubating for 15 min on ice, pelleting at 15,000–21,000g for 15 min at 4°C, resuspending in PBS, centrifuging to pellet insoluble matter, and filtering through 0.45 µm. Heat-inactivated phage were prepared by incubating 1 ml aliquots at 95°C in a water bath for 30 min. Streptomycin-treated mice colonized with Sm^R^ *E. coli* were orally gavaged with 0.2 ml of phage and placed on drinking water containing both streptomycin and carbenicillin.

### Enumeration and culture of *E. coli* from mouse feces

Fecal pellets were collected from individual mice and CFU counts were performed on the same day to determine CFU per gram feces. Briefly, fecal samples (typically 10-40 mg) were weighed on an analytical balance and 250 µl to 500 µl PBS or saline was added. Samples were incubated for 5 min at room temperature and suspended by manual mixing and vortexing. Large particulate matter was pelleted by centrifuging at 100g, ten-fold serial dilutions were made in PBS, and 5 µl of each dilution was spotted on Difco MacConkey agar (BD 212123) supplemented with the appropriate antibiotics, i.e., streptomycin (100 µg/ml) or carbenicillin (50 µg/ml). For qualitative assessment of the fluorescent strains in feces, samples were spotted onto LB supplemented with the appropriate antibiotics. For isolating *E. coli* from fecal samples for genomic or plasmid DNA analysis, the fecal suspension was streaked on agar, and single colonies were further streak-purified.

### Construction of CRISPR-Cas9 phagemid vectors

Cultures were grown in LB or TB media supplemented with the appropriate antibiotics. Plasmid DNA was prepared by QIAprep Spin Miniprep Kit (Qiagen 27106), eluted in TE buffer, and incubated at 60°C for 10 min. Samples were quantified using a NanoDrop One spectrophotometer. The vector pCas9^22^ was digested with BsaI (NEB R0535) and gel extracted with a QIAquick Gel Extraction Kit (Qiagen 28706). Spacers were generated by annealing and phosphorylating the two oligos (PSP116 and PSP117 for GFPT; PSP120 and PSP121 for NT^40^) at 10 µM each in T4 ligation buffer (NEB B0202S) with T4 polynucleotide kinase (NEB M0201S) by incubating at 37°C for 2 h, 95°C for 5 min, and ramping down to 20°C at 5°C/min. The annealed product was diluted 1 in 200 in sterile distilled water and used for directional cloning by ligating (Thermo Scientific FEREL0011) to 60 ng of BsaI-digested, gel extracted pCas9 overnight at room temperature. Ligations were used to transform NEB 5-alpha competent cells (NEB C2987H) and the cloned spacer was verified by Sanger sequencing using primer PSP108. The trailing repeat was later confirmed to lack the starting 5’G, which did not interfere with GFP-targeting function. The 1.8-kb fragment carrying the f1 origin of replication and β-lactamase gene (f1-*bla*) was amplified from pBluescript II with SalI adapters using primers KL215 and KL216 and KOD Hot Start DNA polymerase (Millipore 71842-3). The PCR product was purified using a QIAquick PCR Purification Kit (Qiagen 28104), digested with SalI (Thermo Fisher FD0644), gel extracted, and used to ligate to SalI-digested, FastAP-dephosphorylated (Thermo Fisher FEREF0651) vector. Ligations were used to transform DH5α and clones were screened by restriction digest for both possible insert orientations (A or B) using XbaI (Thermo Scientific FD0684) and one of each orientation was saved for both the GFPT and NT phagemids.

### Preparation of M13 carrying pBluescript II

This protocol was adapted from those to generate phage display libraries^43^. XL1-Blue MRF’ was transformed with pBluescript II (Agilent 212208). An overnight culture of this strain was prepared in 5 ml LB supplemented with tetracycline (5 µg/ml) and carbenicillin (50 µg/ml) and subcultured the following day 1-in-100 into 5 ml 2YT supplemented with the same antibiotics. At an OD_600_ of 0.8, cells were infected with helper phage M13KO7 (NEB N0315S) or VCSM13 (Agilent 200251) at a multiplicity of infection of approximately 10-to-1 for 1 h at 37°C The infected cells were used to seed 2YT supplemented with carbenicillin (100 µg/ml) and kanamycin (25 µg/ml) at 1-in-100, and the culture was grown overnight to produce phage. Cells were pelleted at 10,000g for 15 min, and the supernatant containing phage was transferred. Phage were precipitated by adding 0.2 volumes phage precipitation solution, inverting to mix well, and incubating for 30 min on ice. Phage were pelleted at 15,000g for 15 min at 4°C and the supernatant was discarded. The phage pellet was resuspended in PBS at 1-4% of the culture volume. The resuspension was centrifuged to pellet insoluble material and transferred to a new tube. Glycerol was added to a final concentration of 10-15%. Phage preparations were aliquoted into cryovials and stored at −80°C.

### Preparation of M13 carrying CRISPR-Cas9 phagemids

DH5α(HP4_M13)^44^ was transformed with the GFPT phagemid (pCas9-GFPT-f1A or pCas9-GFPT-f1B) or the NT phagemid (pCas9-GFPT-f1A or pCas9-GFPT-f1B) and plated on LB media containing carbenicillin and kanamycin. Transformants were inoculated into 5 ml 2YT supplemented with 100 µg/ml carbenicillin and 25 µg/ml kanamycin, incubated overnight, used 1-in-100 to seed 250 ml of the same media, and incubated overnight. Cells were pelleted at 10,000g for 15 min, and the supernatant containing phage was transferred. Phage were precipitated by adding 0.2 volumes phage precipitation solution, inverting to mix well, and incubating for 30 min on ice. Phage were pelleted at 20,000g for 20 min at 4°C with slow deceleration. The supernatant was completely removed, phage were resuspended in PBS at 1% of the culture volume, and glycerol was added to a final concentration of 10-15%. The phage solution was centrifuged at 21,000g to pellet insoluble matter, filtered through 0.45 µm, and stored at −80°C.

### Titration of M13 phage carrying phagemid DNA

Phage titer was determined using indicator strain XL1-Blue MRF’ or Sm^R^ W1655 F+. An overnight culture of the indicator strain in LB supplemented with the appropriate antibiotics was subcultured 1-in-100 or 1-in-200 into fresh media and grown to an OD_600_ of 0.8. To estimate titer, serial ten-fold dilutions of the phage preparation were made in PBS, and 10 µl of each dilution was used to infect 90 µl of cells. After incubating at 37°C for 30 min with shaking, 10 µl of the infection mix was spotted onto LB supplemented with carbenicillin. For more accurate titration, 100 µl of phage dilutions were mixed with 900 µl cells in culture tubes, incubated at 37°C for 30 min with shaking, and 100 µl was plated on LB carbenicillin.

Enumeration of viable M13 from mouse feces. Mice were orally gavaged with 6×10 13 M13(pBluescript II) or as negative controls, heat-inactivated phage or PBS. Approximately 100 mg of feces were collected at 0, 3, 6, 9, and 24 h post-gavage, and samples at each timepoint were processed i mmediately. 5 00 μl P BS w as added, samples were incubated for 5 min at room temperature, then suspended by manual mixing and vortexing. Samples were centrifuged at 21,000g for 1 min, the supernatant was transferred to a new tube, and phage titer was determined against i ndicator strain XL1-Blue MRF’ by diluting samples i n PBS, i ncubating with cells, and plating on LB supplemented with carbenicillin. For all dilutions and the undiluted suspension, 10 μl was used to infect 90 μl cells; additionally, for the undiluted suspension, 100 μl was used to infect 900 μl cells to maximize the l imit of detection.

Assay for acid survival. Phage M13(pBluescript II) stored i n PBS was diluted 1-in-100 i n saline. Solutions varying i n pH (1.2, 2, 3, 4, 5, 6, and 7) were prepared by mixing different ratios of 0.2 M sodium phosphate dibasic and 0.1 M citric acid and adjusting with concentrated HCl. 200 μl of each pH solution was transferred to the wells of a microtiter plate, and 10 μl of phage was added containing 1×10 9 M13(pBluescript II). Phage were i ncubated i n the solution, and 10 μl was sampled at 5, 15, and 60 min. Samples were diluted 1-in-100 i n PBS to make acidic samples neutral and phage titer was determined against indicator strain XL1-Blue MRF’ by plating on LB supplemented with carbenicillin. Solution-only controls were assayed simultaneously and cells were plated on LB to confirm viability of the i ndicator strain i n the presence of samples originating from an acidic pH.

### Targeting experiments *in vitro* with M13 CRISPR-Cas9

Overnight cultures of fluorescently marked Sm^R^ W1655 F+ *sfgfp* and *mcherry* were prepared in LB supplemented with streptomycin, subcultured 1 in 200 into fresh media, and grown to an OD_600_ of 0.8. 900 µl cells (approximately 1×10^9^) was transferred to a culture tube, 100 µl phage (approximately 1×10^10^ for f1A vectors and approximately 5×10^10^ for f1B vectors) was added, and the tube was incubated at 37°C for 30 min. The infection culture was transferred to a microfuge tube, cells were pelleted at 21,000g for 1 min, and the supernatant was removed. Cells were washed twice by adding 1 ml PBS, vortexing, pelleting cells, and removing supernatant. Cells were resuspended in 1 ml PBS, and ten-fold serially diluted in PBS. 10 µl of each dilution was spotted onto LB supplemented with carbenicillin and 100 µl was plated on larger plates for isolating single colonies for analysis. Colonies were picked and streak-purified four times to ensure phenotypic homogeneity and clonality.

### Co-culture experiments with *sfgfp* and *mcherry*-marked strains infected with M13 CRISPR-Cas9

Overnight cultures of fluorescently marked Sm^R^ W1655 F+ *sfgfp* and *mcherry* were prepared in LB supplemented with streptomycin. For each culture, three serial ten-fold dilutions were made in PBS, followed by a fourth ten-fold dilution into LB. Equal volumes of each were combined and 5 ml aliquots were transferred to culture tubes. Using a CFU assay, the input was determined to be 6*×*10^6^ CFU of each strain or 1*×*10^7^ CFU total. 10 µl (5*×*10^9^) M13 carrying CRISPR-Cas9 was added, the co-culture was incubated at 37°C for 30 min, and carbenicillin was added to a final concentration of 100 µg/ml. The co-culture was sampled for the t = 0 timepoint and then incubated for 24 h with further sampling every 4 h. At each timepoint, 200 µl was taken; 100 µl was used to assay carbenicillin in the media (see section: Carbenicillin bioassay) and the remaining 100 µl was used for plating as follows. To the 100 µl sample of culture, 900 µl was added and cells were washed by vortexing. Cells were pelleted by centrifuging at 21,000g for 1 min, and 900 µl of the supernatant was removed. To remove residual phage and antibiotic, the wash was repeated once more by adding 900 µl PBS, vortexing, pelleting cells, and removing 900 µl. Cells were resuspended in the remaining 100 µl. Serial ten-fold dilutions were made in PBS and 10 µl of each dilution was spotted onto LB or LB carbenicillin.

### Carbenicillin bioassay

Cultures were sampled over time, cells were pelleted at 21,000g for 1 min, and the supernatant was transferred to a new tube and frozen at −20°C until all timepoints were collected. The supernatants were thawed and assayed using a Kirby-Bauer disk diffusion test. An overnight culture of the indicator organism (*Bacillus subtilis* 168) was diluted in saline to an OD_600_ of 0.1. A cotton swab was dipped into this dilution and spread across LB agar, antibiotic sensitivity disks (Fisher Scientific S70150A) were overlaid using tweezers, and 20 µl of the supernatant was applied to the disk. At the same time, carbenicillin standards were prepared from 1 µg/ml to 100 µg/ml and also applied to discs. Plates were incubated overnight at 37°C and imaged the following morning.

### Flow cytometry

For turbid *in vitro* cultures, samples were diluted 1-in-10,000 in PBS. For mouse fecal pellets, samples were used fresh or thawed from −80°C, and suspended in 500 µl PBS by manual mixing and vortexing. Fecal suspensions were incubated aerobically at 4°C overnight to improve fluorescence signal (**Supplementary** Fig. 18). Samples were vortexed to mix, large particulate matter was pelleted by centrifuging at 100g for 30 seconds, and the sample was diluted 1-in-100 in PBS. Samples were run on a BD LSRFortessa flow cytometer using a 530/30 nm filter for GFP fluorescence and 610/20 nm for mCherry fluorescence, with the following voltages: 750 V for FSC, 400 V for SSC, 700 V for mCherry, and 700-800 V (in vivo) or 650 V (in vitro) for GFP. Flow cytometry data were analyzed in R using packages flowCore (v1.52.1)^45^, Phenoflow (v1.1.2)^46^, and ggcyto (v1.14.0)^47^. Typically, between 10,000 and 100,000 events were collected per sample, and data were rarefied after gating on FSC and SSC. Background events were accounted for on a per-mouse basis. For co-colonization with the *sfgfp*-marked and *mcherry*-marked strains, GFP+ and mCherry+ events from Day −3 (pre-*E. coli*) were used to subtract background at subsequent timepoints. For colonization with the double-marked strain, GFP+ mCherry+ events from Day −5 (pre-*E. coli*) were used to subtract background of double fluorescence at subsequent timepoints, and GFP-mCherry+ events from Day 0 (pre-phage) were used to subtract background of red fluorescence at subsequent timepoints. For exclusion of timepoints due to lack of colonization, the background threshold was calculated as the maximum background observed for that population across all timepoints multiplied by a factor of three. See Code Availability for more information.

### Quick extraction and PCR analysis of genomic DNA from *in vitro* or *in vivo* isolates

Genomic DNA was extracted crudely to use as template for PCR. Briefly, 1.5 ml to 3 ml of culture was transferred to a microfuge tube, cells were pelleted by centrifuging, and the supernatant was discarded. The pellet was frozen, allowed to thaw on ice, resuspended in 100 µl TE, and incubated at 100°C for 15 min in an Eppendorf ThermoMixer. Samples were cooled on ice, cell debris was pelleted by centrifuging at 21,000g for 1 min, the supernatant was transferred to a new tube, and diluted 1-in-100 in TE to use as template DNA. PCR was performed using KOD Hot Start DNA polymerase (Millipore 71842-3) using primers KL207/KL200 for the *sfgfp* gene and primers BAC338F/BAC805R for the 16S rRNA gene^48^.

### Extraction of DNA for hybrid assembly

*E. coli* strains KL68 (W1655 F+ or ATCC 23590), KL114 (W1655 F+ *rpsL*-Sm^R^ *sfgfp*), and KL204 (W1655 F+ *rpsL-*Sm^R^ *sfgfp mcherry*) were cultured in 50 ml LB supplemented with streptomycin. Cells were collected by centrifuging at 6,000g for 10 min at room temperature, washed in 10 ml 10 mM Tris 25 mM EDTA (pH 8.0), and resuspended in 4 ml of the same buffer. 12.5 mg lysozyme (Sigma-Aldrich L6876), 100 µl 5 M NaCl, and 50 µl 10 mg/ml RNase A (Thermo-Fisher EN0531) were added and the mixture was incubated at 37°C for 15 min. To lyse cells, 350 µl 5 M NaCl, 20 µl 20 mg/ml Proteinase K (Ambion AM2546), and 500 µl 10% SDS were added, and the mixture was incubated at 60°C for 1 h with gentle inversions. 2.75 ml of 7.5 M ammonium acetate was added, and the mixture was incubated on ice 20 min to precipitate proteins. Debris was removed by centrifuging 20,000g for 10 min and the supernatant was transferred to a new tube. To extract, an equal volume of chloroform was added and mixed; phases were separated by centrifuging at 2,000g for 10 min, and the aqueous phase was transferred to a new tube. To precipitate the DNA, 1 volume of isopropanol was added, and the tube was inverted until a white precipitate formed. The DNA was pelleted by centrifuging at 2,000g for 10 min and the supernatant was removed. The pellet was washed with 500 µl ice-cold 70% ethanol, allowed to dry, 1 ml TE was added, and the pellet allowed to dissolve overnight at 4°C. To further remove RNA, 250 µl of the genomic prep was transferred to a new tube, 12.5 µl 10 mg/ml RNase A was added, and the mixture was incubated at 37°C for 2 h with mixing every 30 min. To precipitate the DNA, 0.1 volume of 3 M sodium acetate was added followed by 3 volumes of 100% ethanol, and the mixture was inverted until a white precipitate formed. DNA was pelleted by centrifuging at 2,000g for 10 min, the supernatant was removed, the pellet washed with 100 µl 70% ethanol, allowed to dry, and resuspended in 100 µl TE. Samples were quantified by Qubit dsDNA BR Assay and DNA integrity was confirmed by 0.4% agarose gel electrophoresis using GeneRuler High Range DNA Ladder (Thermo-Fisher FERSM1353). DNA was used for both Oxford Nanopore sequencing and Illumina sequencing.

Illumina whole genome sequencing. DNA concentration was quantified using PicoGreen (ThermoFisher). Genomic DNA was normalized to 0.18 ng/ μl f or l ibrary preparation. Nextera XT libraries were constructed i n 384-well plates using a custom, miniaturized version of the standard Nextera XT protocol. Small volume liquid handlers such as the Mosquito HTS (TTP LabTech) and Mantis (Formulatrix) were used to aliquot precise reagent volumes of <1.2 μl to generate a total of 4 μl per library. Libraries were normalized and 1.2 μl of each normalized library was pooled and sequenced on the Illumina NextSeq or MiSeq platform using 2×146 bp configurations. 12 bp unique dual indices were used to avoid index hopping, a phenomenon known to occur on ExAmp based Illumina technologies. See Data Availability for more information.

### Oxford Nanopore sequencing and hybrid Nanopore/Illumina assembly

PCR-free long read libraries were prepared using the Ligation Sequencing Kit (SQK-LSK109), multiplexed using the Native Barcoding Kit (EXP-NBD114), and sequenced on the MinION platform using flow cell version MIN106 (Oxford Nanopore Technologies). Basecalling of MinION raw signals was done using Guppy (v2.2.2, Oxford Nanopore Technologies). Reads were demultiplexed with qcat (v1.1.0, Oxford Nanopore Technologies). Quality control was achieved using porechop (v0.2.3 <u>s</u>eqan2.1.1) (https://github.com/rrwick/Porechop) using the discard middle option. Reads were filtered using NanoFilt (v2.6.0)^49^ with the following parameters: minimum average read quality score of 10 (-q 10) and minimum read length of 100 (-l 100). Illumina reads were quality filtered using fastp (v0.20.1)^50^ with the following parameters: cut front, cut tail, cut <u>w</u>indow <u>s</u>ize 4, cut mean <u>q</u>uality 20, length required 60. Filtered MinION and Illumina reads were then provided to Unicycler (v0.4.8)^51^ for hybrid assembly; default parameters were used unless otherwise noted. See Data Availability for more information.

### Analysis of isolates after *in vitro* or *in vivo* M13-mediated delivery of phagemid

Isolates were cultured on LB or Difco MacConkey agar plates supplemented with carbenicillin or both carbenicillin and streptomycin. Isolates from *in vitro* GFP-targeting experiments were streak purified 4 times on agar to ensure clonality. Isolates from *in vivo* experiments were obtained by suspending a fecal pellet in 500 µl PBS, streaking the suspension onto agar, followed by streak purification of single colonies. For DNA extraction, single colonies were inoculated into LB or TB supplemented with the appropriate antibiotics. For analysis of the phagemid, plasmid DNA was extracted using a QIAprep Spin Miniprep Kit (Qiagen 27106), eluted in TE buffer, and incubated at 60°C for 10 min. DNA was quantified using a NanoDrop One spectrophotometer and 200-600 ng was digested with FastDigest restriction enzymes (KpnI, Thermo Scientific FD0524; XbaI, Thermo Scientific FD0684) for 10 min at 37°C followed by gel electrophoresis. Spacer sequences on phagemids were confirmed by Sanger sequencing using primer PSP108. For genome sequencing, genomic DNA was either extracted using a DNeasy Blood & Tissue Kit (Qiagen 69506) or an in-house protocol. Briefly, isolates were cultured in 3 ml TB supplemented with streptomycin and carbenicillin. Cells were pelleted and resuspended in 460 ul of freshly prepared buffer [per sample: 400 ul 10 mM Tris (pH 8.0) 25 mM ETDA, 50 µl 5 M NaCl, and 10 µl 10 mg/ml RNase A (Thermo-Fisher EN0531)]. 50 µl 10% SDS was added, mixed well, and samples were incubated at 60°C for 1 h with periodic inversions. 260 µl of 7.5 M ammonium acetate was added, and the mixture was incubated on ice 20 min to precipitate proteins. Precipitate was removed by centrifuging 21,000g for 5 min and the supernatant was transferred to a new tube. To extract, an equal volume of chloroform was added and mixed; phases were separated by centrifuging at 21,000g for 2.5 min. The aqueous phase was transferred to a new tube, centrifuged at 21,000g for 2.5 min, and 500 ul was transferred to a new tube. To precipitate the DNA, 500 µl isopropanol was added, and the tube was inverted until a white precipitate formed. Using a pipette tip, the clump was transferred to a new tube, washed with 100 µl cold 70% ethanol, and allowed to dry. 50 µl TE was added and the pellet was allowed to dissolve at 4°C overnight. DNA integrity was confirmed by gel electrophoresis and used for Illumina whole genome sequencing (see section: Illumina whole genome sequencing). Sequence reads were quality filtered using fastp (v0.20.1)^50^ and reads were aligned using bowtie2 (v2.3.5.1)^52^ to reference genomes and phagemid sequences; complete reference genomes were generated using hybrid assembly (see section: Oxford Nanopore sequencing and hybrid Nanopore/Illumina assembly). For Carb^R^ isolates obtained after delivery of the pBluescript II phagemid, reads were simultaneously aligned to the genome of strain KL68 (W1655 F+) and the pBluescript II sequence (NCBI accession X52329.1). For Carb^R^ isolates obtained after delivery of CRISPR-Cas9 phagemids, reads were simultaneously aligned to the genome of the strain used for *in vitro* (KL114; *sfgfp*) or *in vivo* (KL204; *sfgfp mcherry*) experiments and the sequence of the delivered phagemid. For isolates from GFP-targeting experiments, deletions were visualized by using samtools (v1.9)^53^ to filter multi-mapping and low-quality read alignments with MAPQ<2 (view -q 2), and depth was calculated using a sliding window of 20; breseq (v0.35.4)^54^ was used to assess deletion size. See Data Availability for more information.

## Data availability

All raw sequence data have been deposited at the NCBI Sequence Read Archive under BioProject PRJNA642411. For 16S rRNA gene sequencing, Illumina sequence data have been deposited with accession numbers SRR12118792 to SRR12118959. For sequencing of isolates from mouse fecal samples after delivery of pBluescript II, Illumina sequence data deposited with accession numbers SRR14278062 to SRR14278073. For Nanopore/Illumina hybrid assembly of reference strain KL68 (W1655 F+ or ATCC 23590) as well as Sm^R^ fluorescent derivatives KL114 (*sfgfp*) and KL204 (*sfgfp mcherry*), Illumina sequence data have been deposited with accession numbers SRR14296642 to SRR14296644 and Oxford Nanopore MinION data with accession numbers SRR14297452 to SRR14297454. For sequencing of isolates after targeting with phage-delivered CRISPR-Cas9, Illumina sequence data deposited with accession numbers SRR14289086 to SRR14289109. Supporting data have also have been made publicly available on Github: https://github.com/turnbaughlab/2021_Lam_M13_CRISPRCas9

## Code availability

Bash scripts and R Markdown documents for flow cytometry, 16S rRNA gene sequencing, and genomic deletion analyses have been made publicly available on GitHub: https://github.com/turnbaughlab/2021_Lam_M13_CRISPRCas9

## Acknowledgments

We thank the UCSF Gnotobiotics Core staff (Jessie Turnbaugh, Kimberly Ly, and Jolie Ma) for animal care support and assistance with learning animal procedures. We are grateful to Daryll Gempis, Bernarda Lopez, and Ernesto Valencia of the UCSF G.W. Hooper Foundation for laboratory and administrative support. We thank Katja Engel (University of Waterloo) for help in translating the original publication on the isolation of M13 written in German. We are grateful to Antoine Vigouroux (Institut Pasteur) for generously sharing MG1655 derivatives carrying *sfgfp* and *mcherry* marker genes. We thank the Chan-Zuckerberg BioHub for sequencing through the Microbiome Initiative. We are grateful to Joseph Bondy-Denomy and Oren Rosenberg (UCSF) for constructive criticism of the manuscript. The graphic of a laboratory mouse was adapted from a Wikimedia Commons graphic distributed under CC-BY-SA-4.0 by Gwilz. KNL and PS were both supported by a postdoctoral fellowship from the Canadian Institutes of Health Research (CIHR), JEB by a postdoctoral fellowship from the Natural Sciences and Engineering Research Council of Canada (NSERC), MA by an F32 fellowship from the National Institutes of Health (F32AI147456-01), and PSP by a scholarship from the UCSF Discovery Fellows Program. This work was supported by the National Institutes of Health (PJT, R01HL122593, R01AT011117; RRN, K08AR073930). PJT held an Investigators in the Pathogenesis of Infectious Disease Award from the Burroughs Wellcome Fund, was a Chan Zuckerberg Biohub investigator, and was a Nadia’s Gift Foundation Innovator supported, in part, by the Damon Runyon Cancer Research Foundation (DRR-42-16) and the Searle Scholars Program (SSP-2016-1352).

## Author contributions

KNL and PJT conceived the ideas. KNL supervised laboratory work and analyzed the data. KNL, PS, PSP, and PJT designed the experiments. KNL and PSP constructed plasmids. KNL constructed *E. coli* strains with assistance from PS. KNL performed phage and animal experiments with assistance from PSP. KL prepared samples and PS performed antibiotic assays. KNL, PSP and MN prepared samples, FBY performed 16S rRNA gene sequencing, and KNL and JEB analyzed the data. MA performed flow cytometry and KNL analyzed the data. KNL analyzed plasmid DNA from mouse fecal isolates with assistance from MN. KNL prepared samples, FBY and AMW performed Nanopore and Illumina whole genome sequencing, RRN performed the hybrid assembly of reference genomes, and KNL analyzed the genomic deletion data. PJT provided reagents and materials. KNL made the figures. KNL and PJT wrote the manuscript; PS, PSP, MA, JEB, and FBY assisted with editing.

## Additional information

Correspondence and requests for materials should be addressed to PJT.

## SUPPLEMENTARY INFORMATION

### Supplementary Figures

**Supplementary Figure 1.**
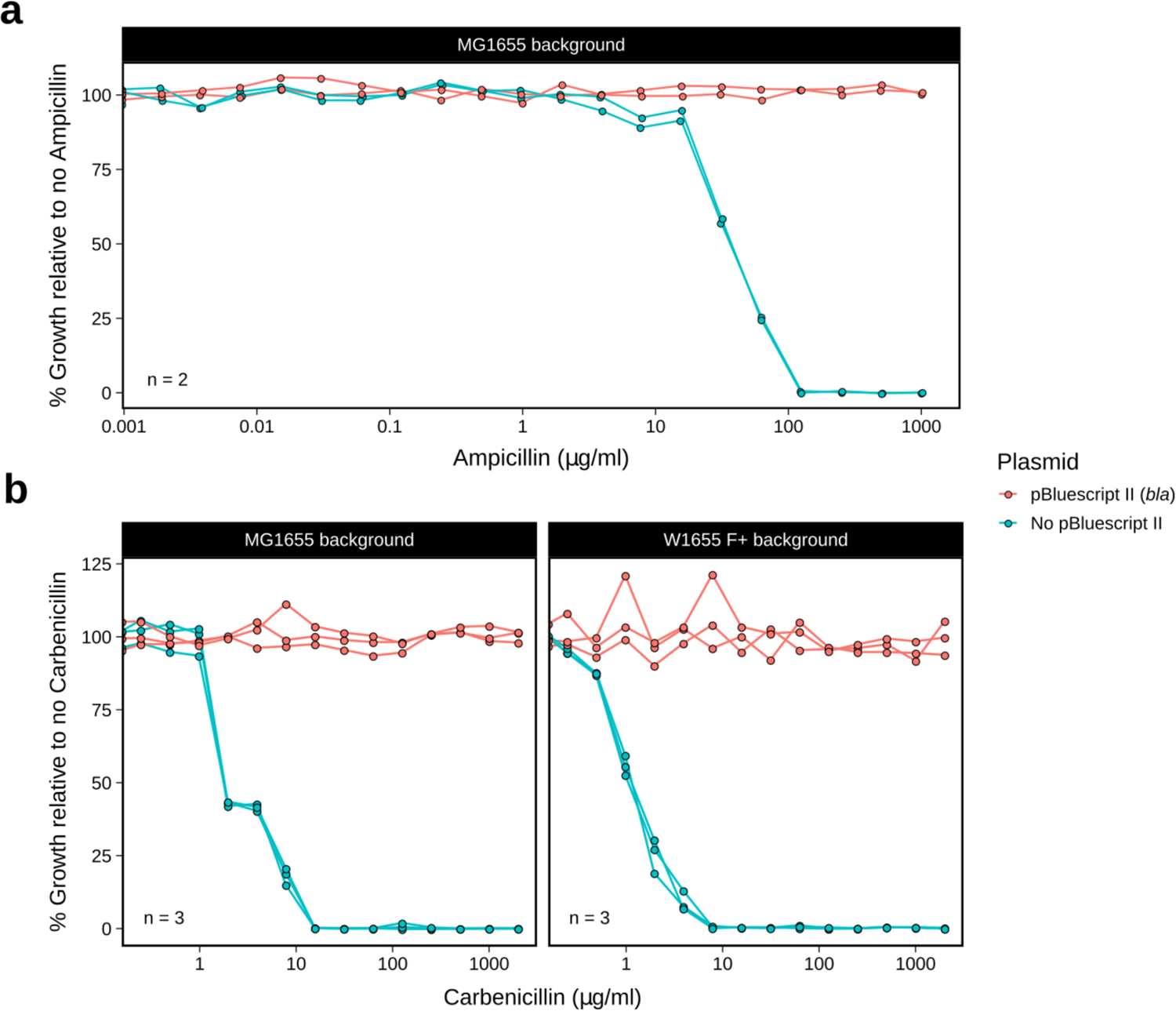
The plasmid pBluescript II confers resistance to beta-lactam antibiotics exceeding 1 mg/ml. (a) Minimum inhibitory concentration (MIC) assay for ampicillin in the *E. coli* MG1655 background. Harbouring pBluescript II, the strain exhibits an MIC of >1 mg/ml; in the absence of the plasmid, the MIC is approximately 100 µg/ml (n=2 biological replicates). **(b)** pBluescript II confers resistance to carbenicillin exceeding 2 mg/ml in both the *E. coli* MG1655 and W1655 F+ backgrounds; in the absence of the plasmid, the same strains have an MIC of approximately 10 µg/ml (n=3 biological replicates per strain). *bla*, beta-lactamase gene.

**Supplementary Figure 2.**
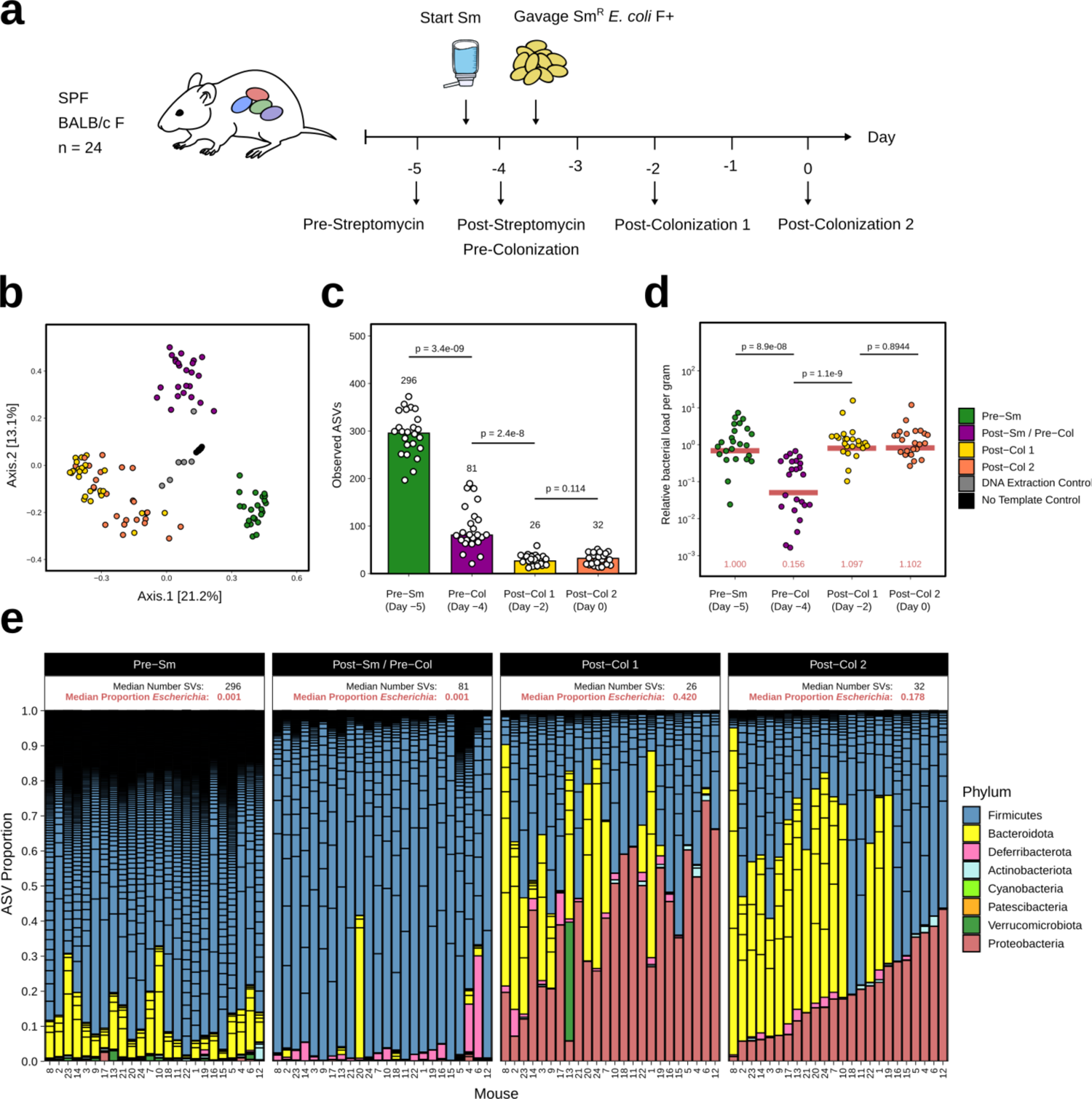
Streptomycin treatment reduces bacterial diversity and allows Sm^R^ *E. coli* to colonize at a high proportion in conventionally raised mice. (a) Typical timeline for a mouse experiment using water containing streptomycin (Sm) to allow colonization by Sm^R^ *E. coli.* Fecal samples of individually caged mice (n=24) were collected before streptomycin (Day −5), after streptomycin but before gavage of *E. coli* (Day −4), and two timepoints after *E. coli* (Day −2 and 0). (b) Principle coordinate analysis using Bray-Curtis dissimilarity indicates that fecal samples from Pre-Sm, Post-Sm/Pre-Colonization (Pre-Col), and Post-Colonization (Post-Col) are distinct. (c) Number of observed 16S rRNA gene amplicon sequence variants (ASVs) was lower in Post-Sm relative to Pre-Sm timepoint, and lower still in both Post-Col timepoints than Pre-Sm and Post-Sm/Pre-Col. Numbers above bars indicate median. (d) The bacterial load in fecal samples (based on qPCR of 16S rRNA gene) transiently decreased during the course of treatment. Red numbers and horizontal bar indicate median for each timepoint. (e) Proportion of individual ASVs detected in each mouse at the different timepoints, coloured by phylum. Where ASV count is high, *e.g.*, in Pre-Sm timepoint, stacked bars appear to fade to black due to a large number of low abundance ASVs. For each group, the median number of ASVs as well as the median proportion of ASVs classified as *Escherichia-Shigella* is indicated. *p*-value, Mann-Whitney test.

**Supplementary Figure 3.**
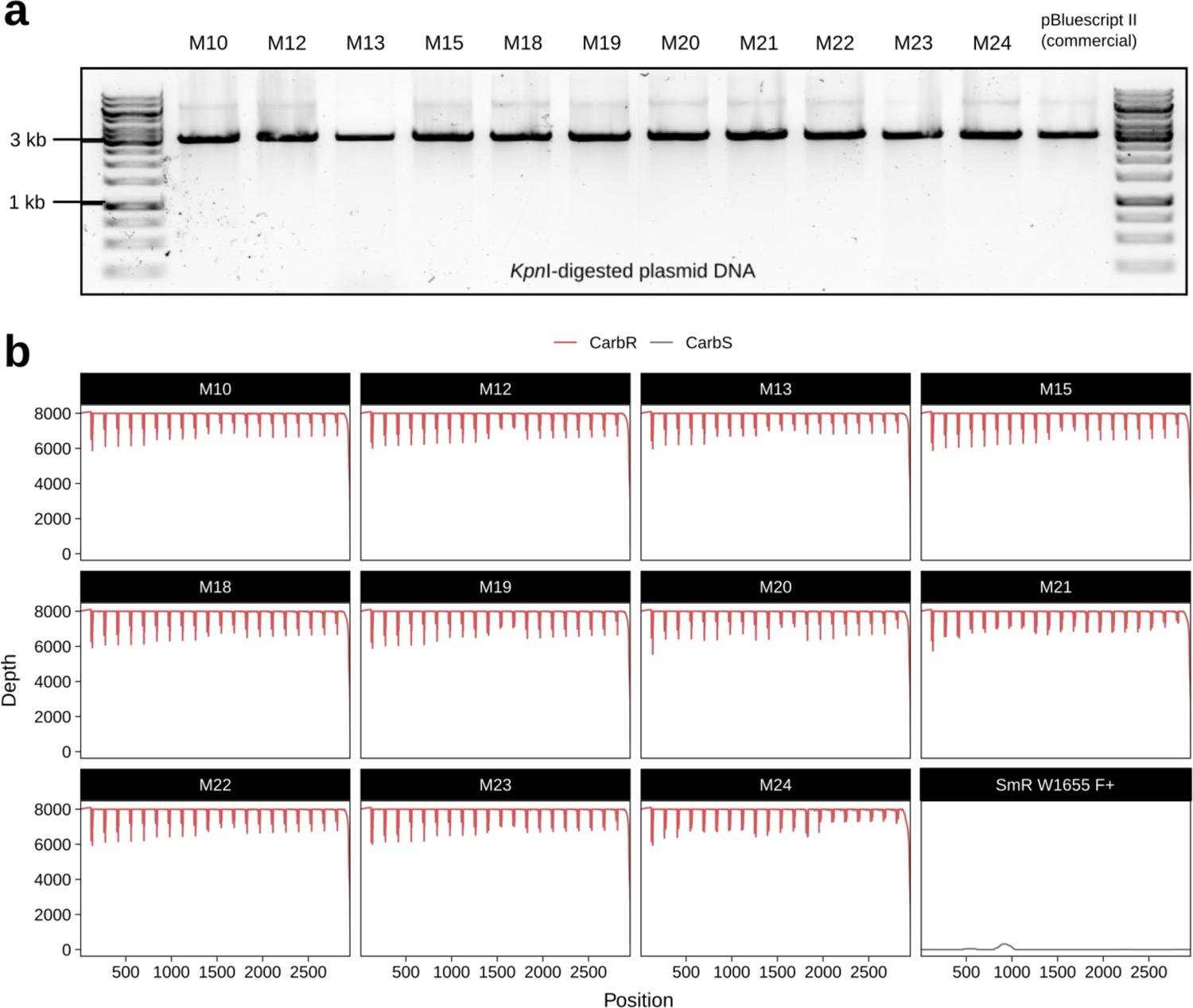
Successful transfer of pBluescript II plasmid from M13 phage to *E. coli* cells *in vivo*. **(a)** Diagnostic digest of extracted plasmid DNA from fecal isolates is consistent with pBluescript II. Plasmid DNA was recovered from carbenicillin-resistant (Carb^R^) colonies isolated from the feces of the 11 mice that were successfully colonized during treatment with carbenicillin in the water (**Fig. 1d**); DNA was digested with restriction enzyme KpnI for comparison to linearized 3-kb pBluescript II. **(b)** Genome sequencing results from Carb^R^ isolates confirms presence of pBluescript II. Genomic DNA was extracted from these isolates and sequenced; reads were mapped to the reference W1655 F+ genome and pBluescript II; read depth for the 2961-bp plasmid is shown. As a negative control, the Sm^R^ W1655 F+ strain used to colonize these mice at the start of the experiment [prior to treatment with M13(pBluescript II)] was sequenced in the same batch.

**Supplementary Figure 4.**
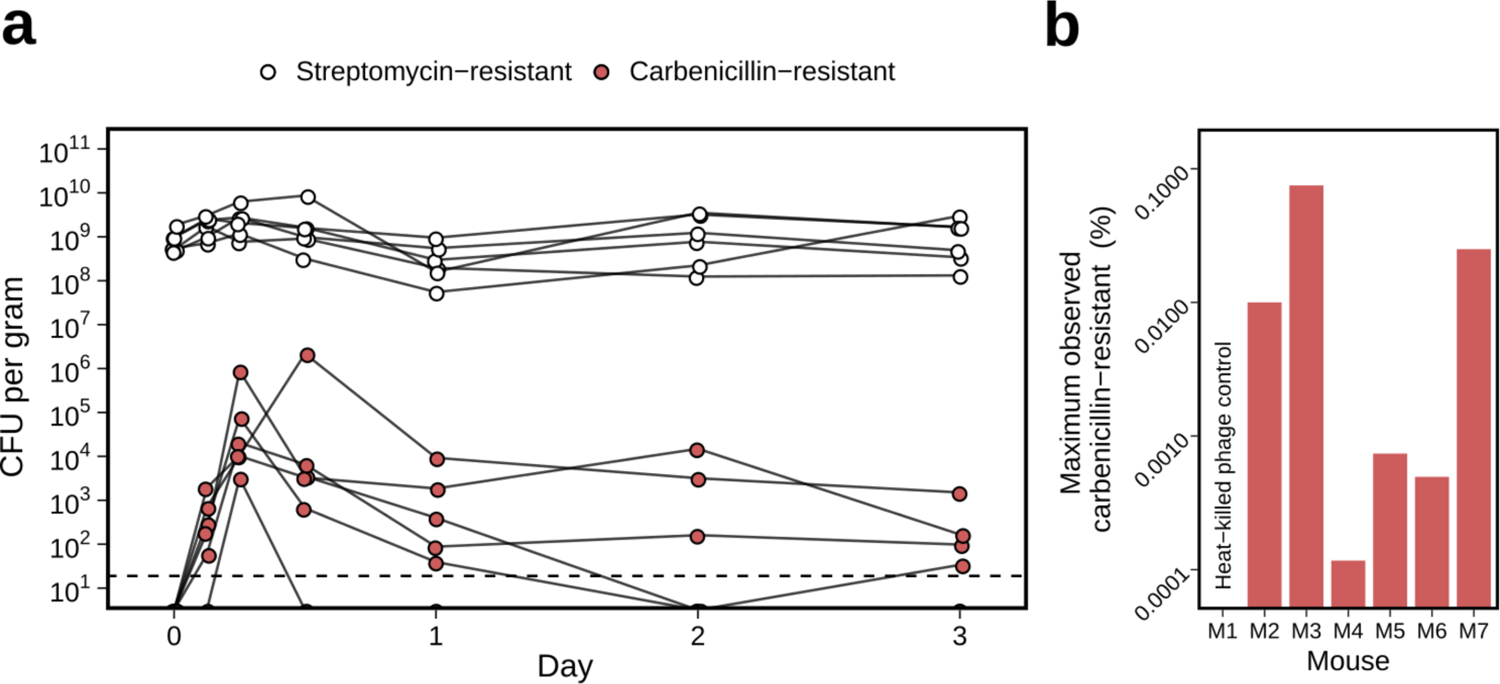
Low frequency of carbenicillin-resistant *E. coli* following treatment with M13(pBluescript II) in the absence of carbenicillin selection. (a) Sm^R^ *E. coli* W1655 F+ was introduced and maintained in streptomycin-treated mice (n=6). Oral gavage of 10^13^ M13(pBluescript II) was performed on Day 0, but contrary to **Fig. 1c**, carbenicillin was not added to the drinking water to select for phage infection. Sm^R^ and Carb^R^ CFU were enumerated up to 3 days post-gavage. Sm^R^ CFU indicative of total *E. coli* remained steady while Carb^R^ CFU indicative of phage-infected cells were much lower in number, peaked between 6-12 h after gavage, and decreased over time. Dashed line indicates limit of detection. (b) The maximum observed percentage of Carb^R^ CFU (phage-infected) over Sm^R^ CFU (total) was approximately 0.1%. For each of the six mice in panel a, the maximum percentage of Carb^R^/Sm^R^ was calculated, with values ranging over 3 orders of magnitude between ∼0.0001% and ∼0.1%. Time-series fecal samples from a mouse orally gavaged with heat-killed phage was also assayed as a negative control.

**Supplementary Figure 5.**
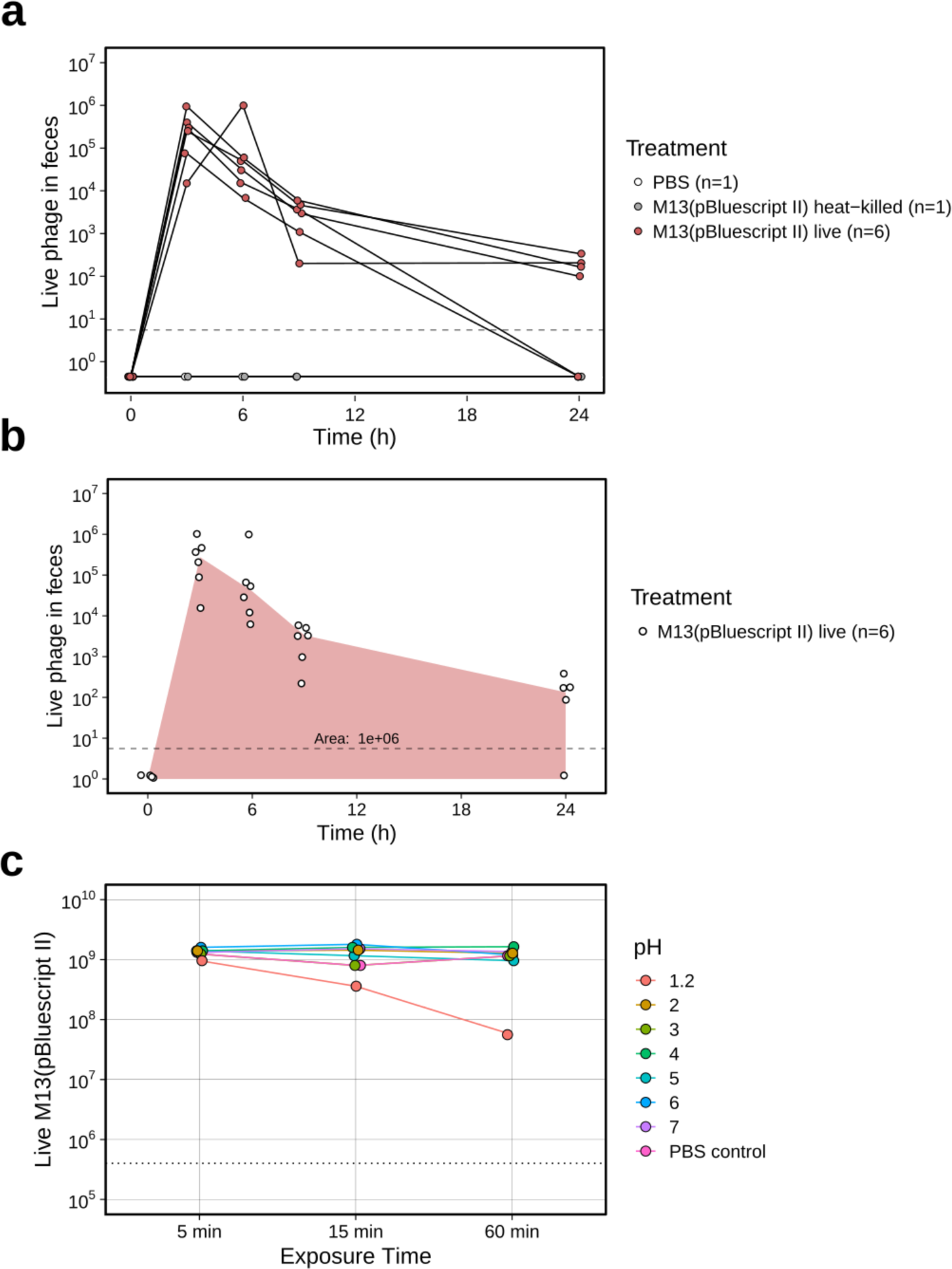
Low recovery of M13 bacteriophage following GI transit despite high acid tolerance. (a) Enumeration of phage M13(pBluescript II) in feces of conventionally-raised mice after oral gavage. Mice were treated with 10^13^ M13(pBluescript II) (n=6) or as negative controls, heat-killed M13(pBluescript II) or PBS. Live phage in fecal samples were assayed using indicator strain XL1-Blue MRF’ at t = 0, 3, 6, 9, and 24 h post-gavage. **(b)** Using the median live phage output in the feces at each timepoint for the six mice gavaged with live M13(pBluescript II), the area under-the-curve from 0 to 24 hours is 1*×*10^6^. **(c)** Phage M13 displays resistance to acidic conditions as extreme as pH 2. 10^9^ M13(pBluescript II) were incubated in pH solutions 1.2 to 7, and sampled over the course of 60 min to assay for viability. Dashed line indicates limit of detection.

**Supplementary Figure 6.**
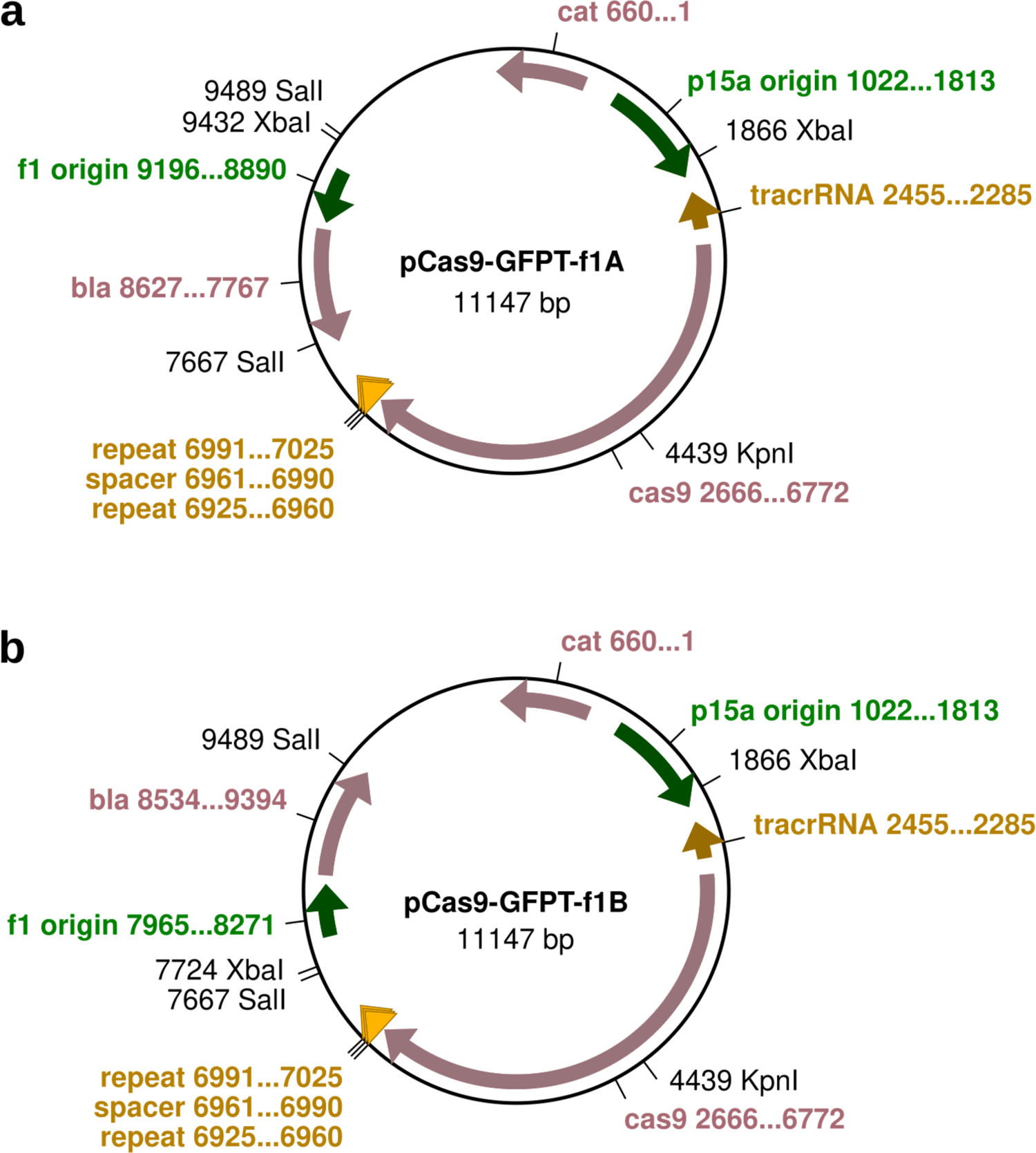
GFP-targeting (GFPT) CRISPR-Cas9 phagemids. The non-targeting (NT) versions of these vectors (not shown here) are identical to the GFPT vectors except in the spacer sequence. The f1-*bla* fragment was cloned as a SalI fragment in both possible orientations for either strand of DNA to be packaged into M13 phage. **(a)** The first orientation is designated f1A. **(b)** The second orientation is designated f1B. *cat*, chloramphenicol acetyltransferase (Cm^R^); *bla*, beta-lactamase (Carb^R^).

**Supplementary Figure 7.**
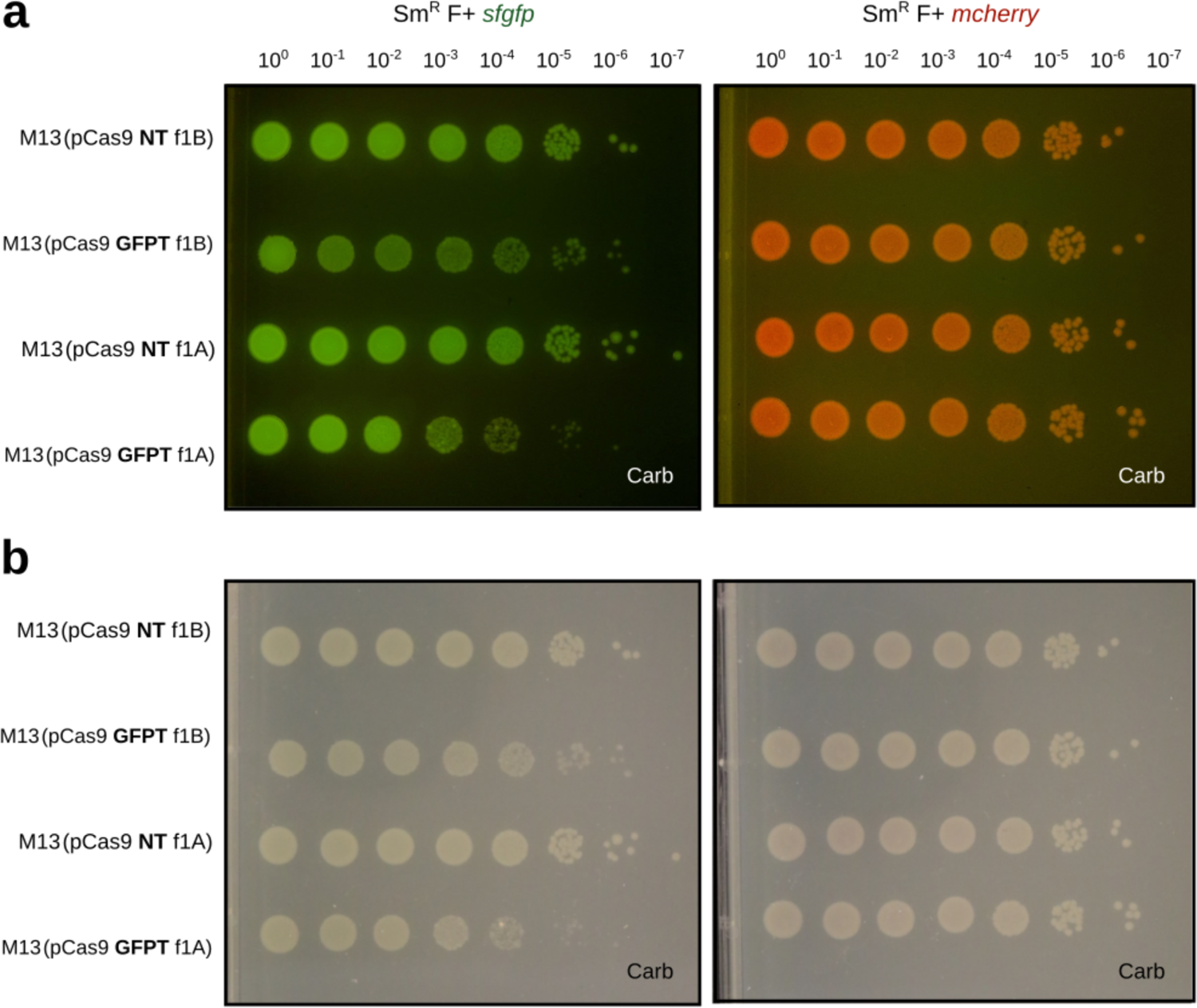
GFP-marked *E. coli* exhibits impaired colony growth after infection with GFPT-M13 *in vitro*. NT-M13 or GFPT-M13 were used to infect Sm^R^ W1655 F+ *sfgfp* or Sm^R^ W1655 F+ *mcherry* (negative control). **(a)** Growth impairment of the GFP-marked strain under GFPT conditions was evident under blue light. **(b)** Impaired colonies exhibited a translucent quality that was more pronounced under white light.

**Supplementary Figure 8.**
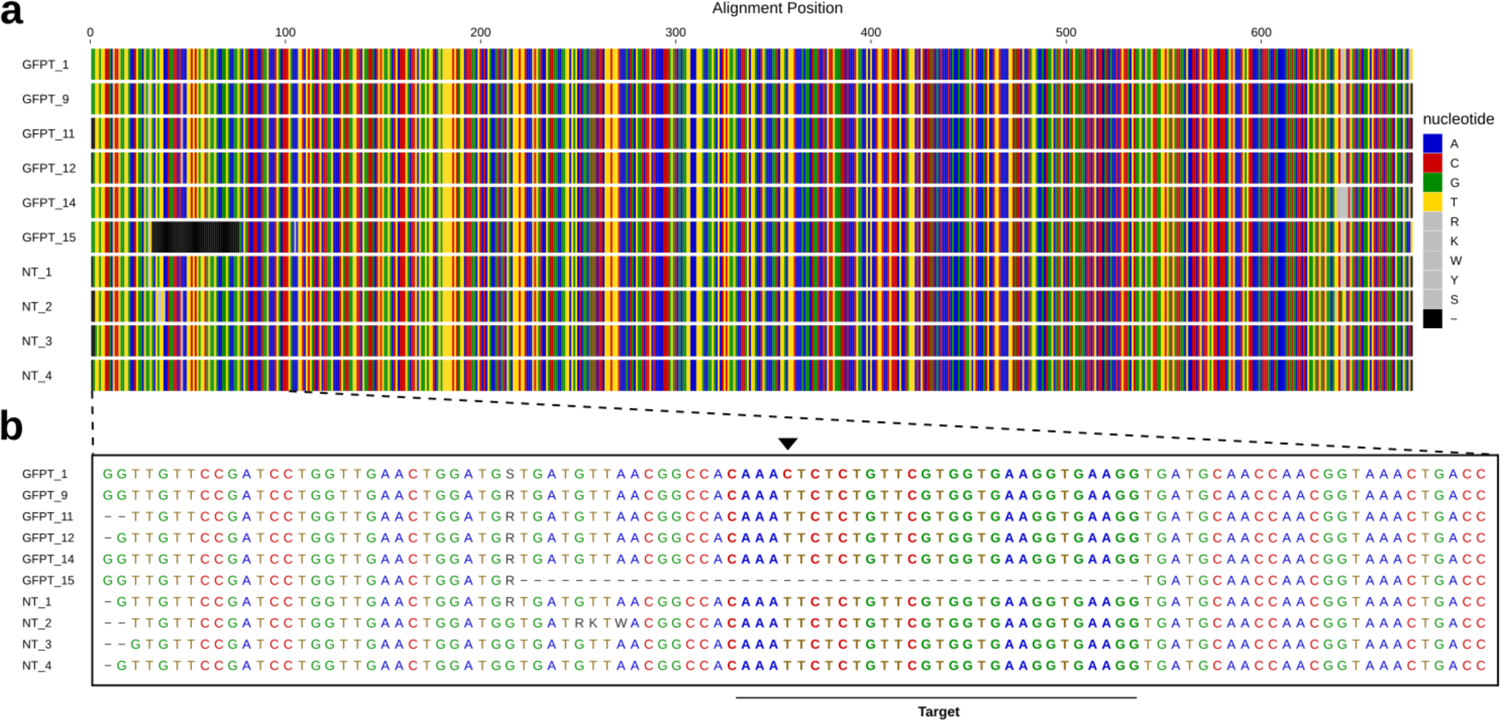
Escape from GFP-targeting CRISPR-Cas9 by mutation in the target sequence. (a) Sanger sequencing of *sfgfp* PCR amplicons from streak-purified clones after treatment with GFPT-M13 or NT-M13 *in vitro* (**Fig. 2c**) confirmed the partial loss observed for clone GFPT 15 by gel electrophoresis. **(b)** Pullout showing the lost region of the *sfgfp* coding sequence from clone GFPT 15 encompasses the CRISPR-Cas9 target site. Closer examination of clone GFPT 1 revealed the presence of a single nucleotide change in the target site (black arrow) that allowed this clone to escape targeting yet remain fluorescent.

**Supplementary Figure 9.**
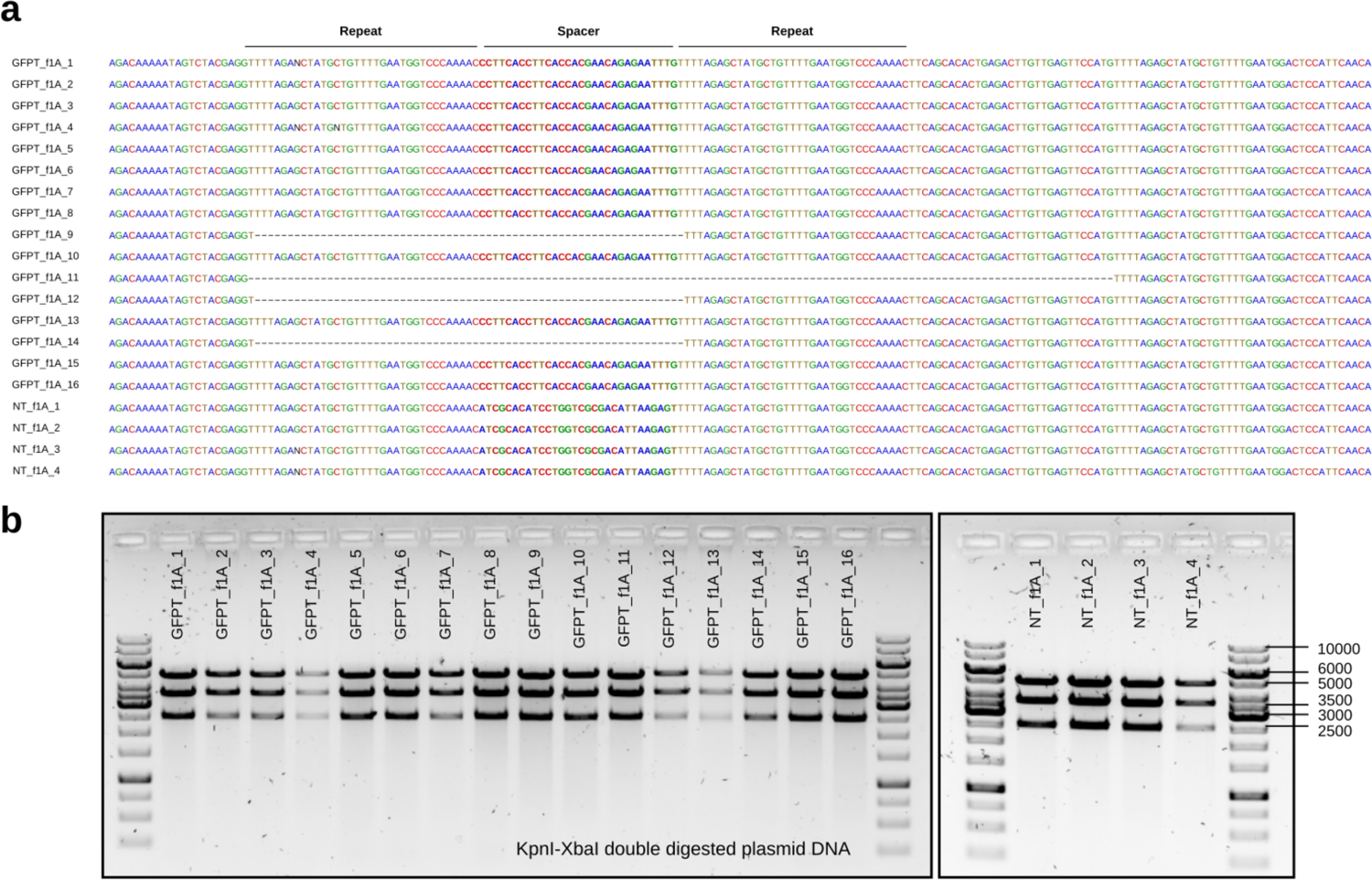
Spacer loss from GFP-targeting CRISPR-Cas9 phagemids. **(a)** Sanger sequencing to check for spacer presence in phagemid DNA isolated from clones after treatment with GFPT-M13 or NT-M13 *in vitro*. All 4 clones isolated after infection with NT-M13 retained the spacer. Of 16 clones isolated after infection with GFPT-M13, 4 had lost the spacer (clones 9, 11, 12, and 14). **(b)** Diagnostic digest of plasmid DNA isolated from clones using KpnI and XbaI revealed phagemid of the expected size. Expected fragments: 4993, 3581, and 2573 bp.

**Supplementary Figure 10.**
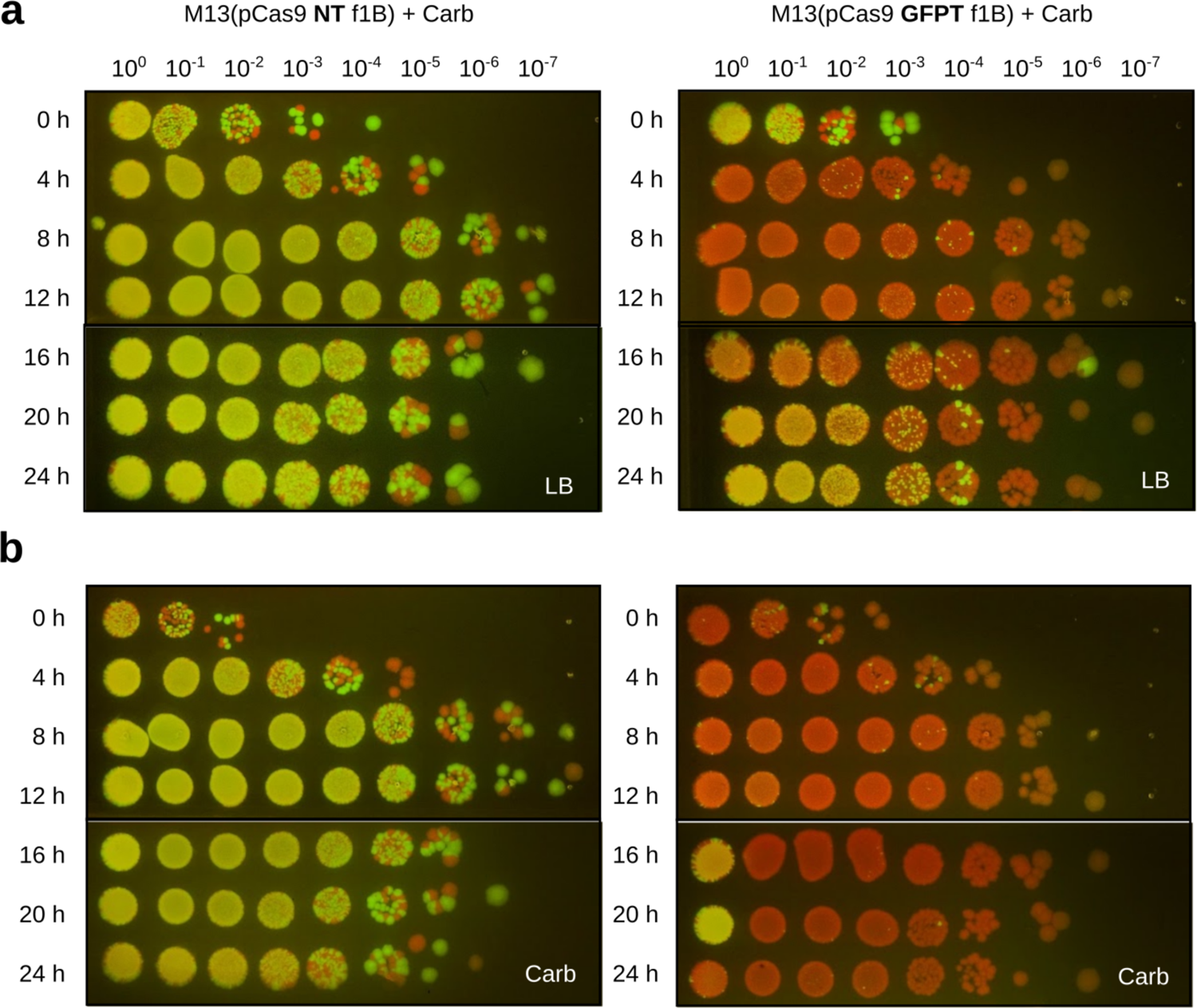
Recovery of GFP+ cells at later timepoints from *in vitro* co-cultures after infection with GFPT-M13 is likely due to lack of selection for the CRISPR-Cas9 phagemid. (a) On non-selective media, GFP fluorescent colonies are detected at later timepoints of the co-culture infected with GFPT-M13. (b) Lack of GFP fluorescent colonies after testing the same co-culture on media with carbenicillin indicates that those GFP+ colonies at later timepoints derive from cells that are Carb^S^ and suggests that they do not harbour the CRISPR-Cas9 phagemid.

**Supplementary Figure 11.**
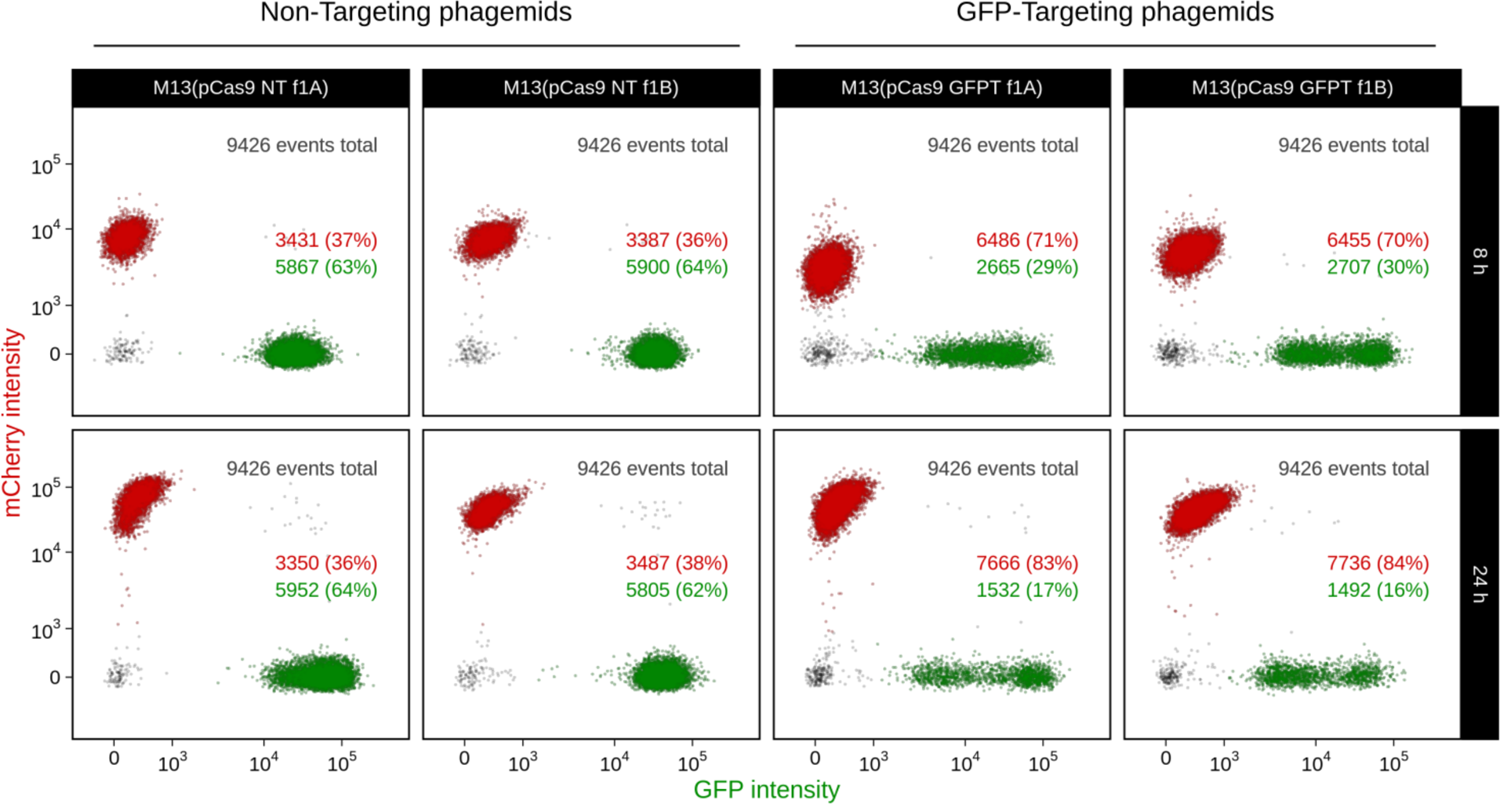
The GFP+ strain shows a further decrease in relative abundance at 24 h after treatment with GFPT-M13 when compared to the 8 h timepoint. Co-cultures of GFP-marked and mCherry-marked *E. coli* F+ were infected with NT-M13 or GFPT-M13, carbenicillin was added to select for phage infection, and fluorescent populations were assayed by flow cytometry. The relative abundance of GFP+ events is decreased in GFPT conditions at 8 h and further decreased by 24 h. Non-targeting phagemids are pCas9-NT-f1A and pCas9-NT-f1B; GFP-targeting phagemids are pCas9-GFPT-f1A and pCas9-GFPT-f1B.

**Supplementary Figure 12.**
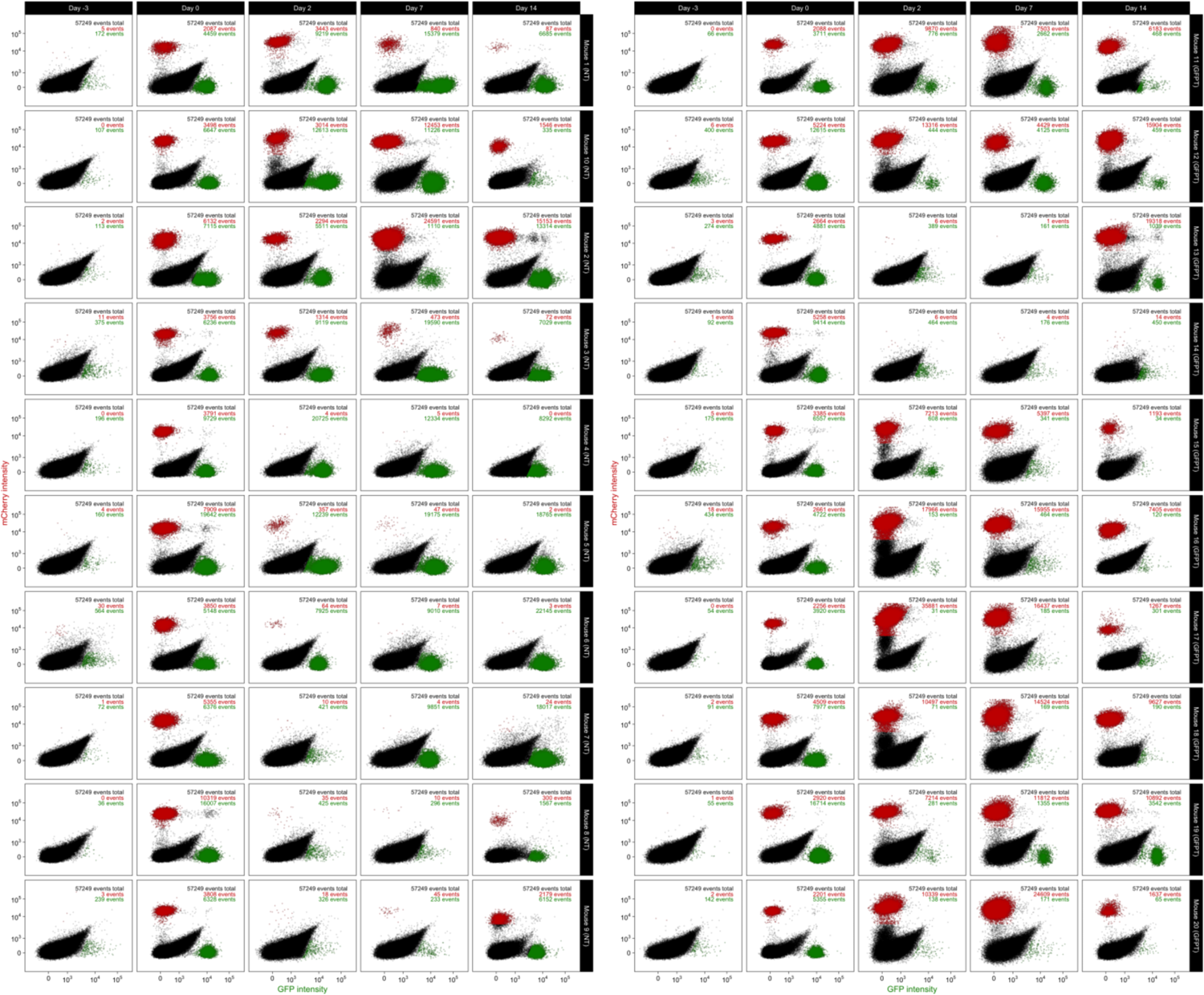
Flow cytometry plots of fecal samples for all mice at all timepoints during in vivo targeting of GFP-marked *E. coli* in competition with mCherry-marked *E. coli*. Mice (n = 10 per group) were given either NT-M13 (left) or GFPT-M13 (right). Day −3, before colonization by *E. coli*; Day 0, after colonization by both GFP+ and mCherry+ strains; Day −2, post phage and carbenicillin treatment; Day 7, one week post-phage and carbenicillin; Day −14, one week after removing carbenicillin from drinking water.

**Supplementary Figure 13.**
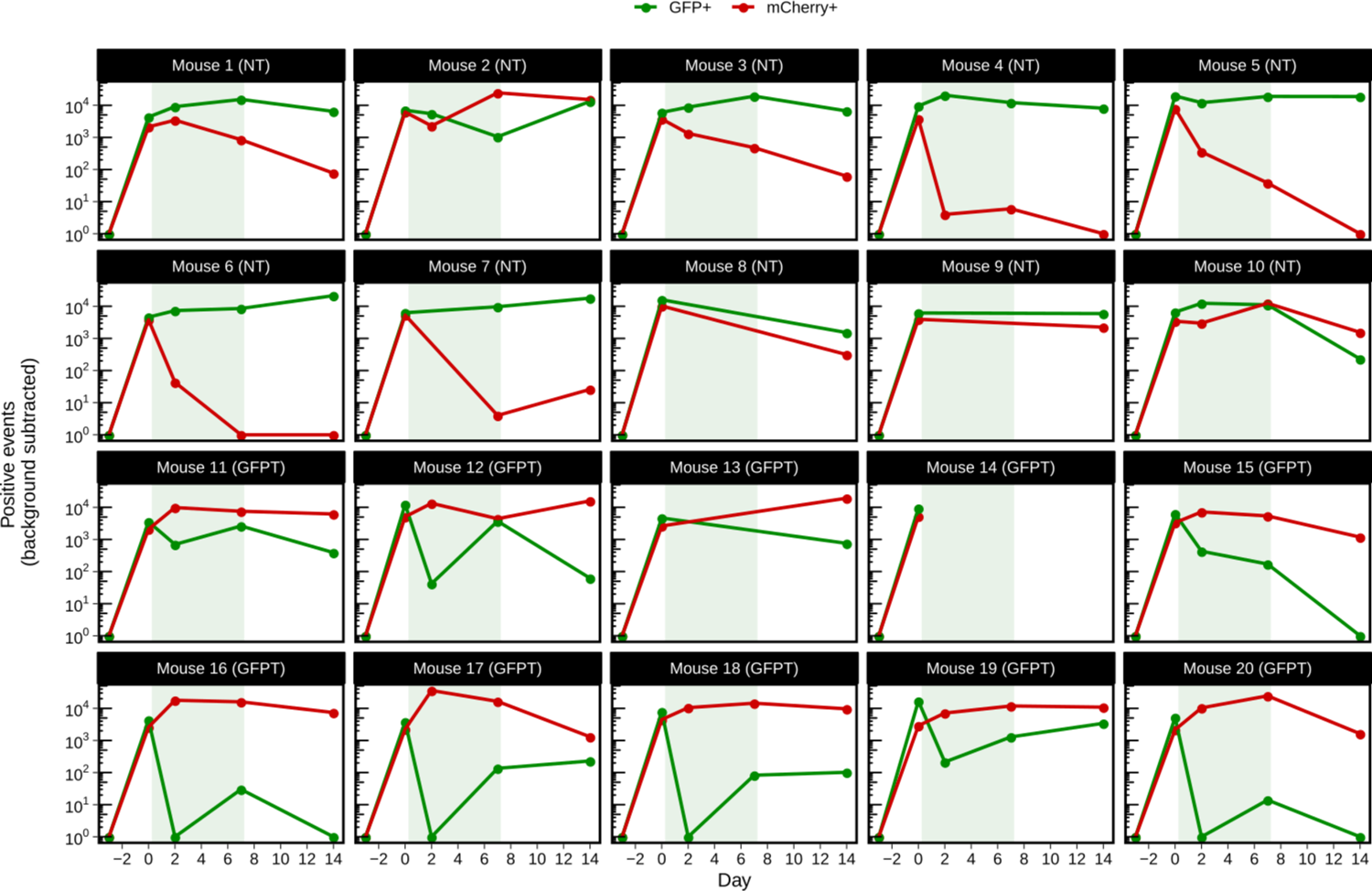
GFP+ and mCherry+ events in fecal samples over time for individual mice during in vivo targeting of GFP-marked *E. coli* in competition with mCherry-marked *E. coli*. Mice were treated with either NT-M13 (M1 to M10) or GFPT-M13 (M11 to M20). For each mouse, the number of positive events by flow cytometry on Day −3 (before *E. coli* colonization) was used to subtract background for all subsequent timepoints. Shaded green area indicates duration of carbenicillin treatment. Timepoints were excluded when both mCherry+ and GFP+ events were below background thresholds.

**Supplementary Figure 14.**
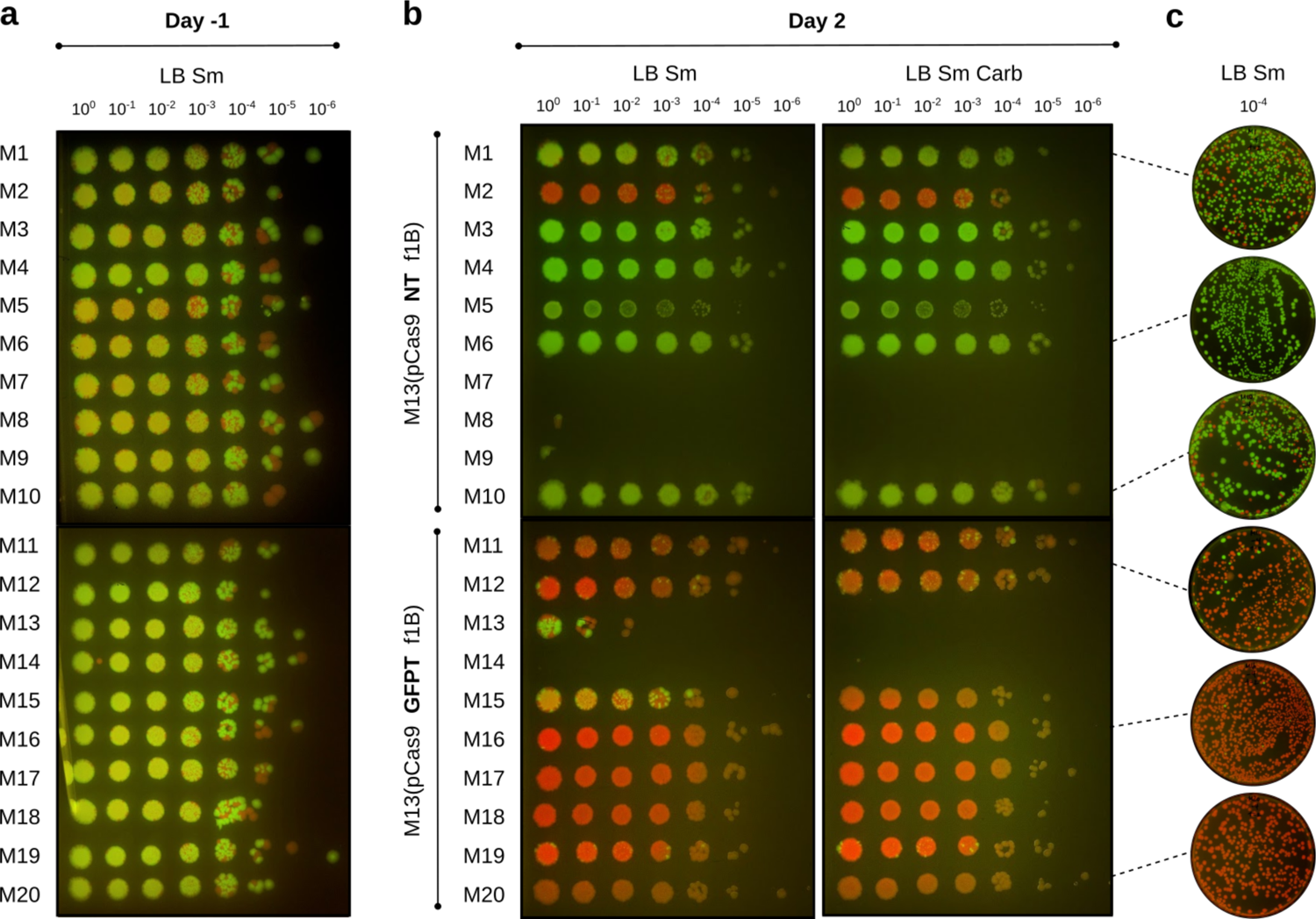
Culturing from fecal samples of mice co-colonized with GFP-marked and mCherry-marked strains before and after treatment with phage confirms depletion of GFP+ strain under GFPT conditions on Day 2. (a) At Day −1, colonization by both the GFP+ and mCherry+ *E. coli* was confirmed in fecal samples of all mice by culturing on LB with streptomycin. (b) After treating with NT-M13 (M1 to M10) or GFPT-M13 (M11 to M20) and carbenicillin to select for phage infection, culture of *E. coli* on LB streptomycin (Sm) from fecal samples on Day 2 of GFPT mice exhibit decreased GFP fluorescence. Culturing from the same samples on LB with both streptomycin and carbenicillin (Carb) suggests that for some mice, fluorescent colonies arising on LB streptomycin are Carb^S^, *i.e.*, that they do not carry the CRISPR-Cas9 phagemid. Lack of fluorescent *E. coli* in fecal samples indicates eradication by carbenicillin where phage infection leading to colonization by Carb^R^ *E. coli* has not occurred. (c) Day 2 fecal suspensions from a subset of the mice (M1, M6, M10 for NT; M11, M16, M20 for GFPT) were cultured on larger plates for confirmation.

**Supplementary Figure 15.**
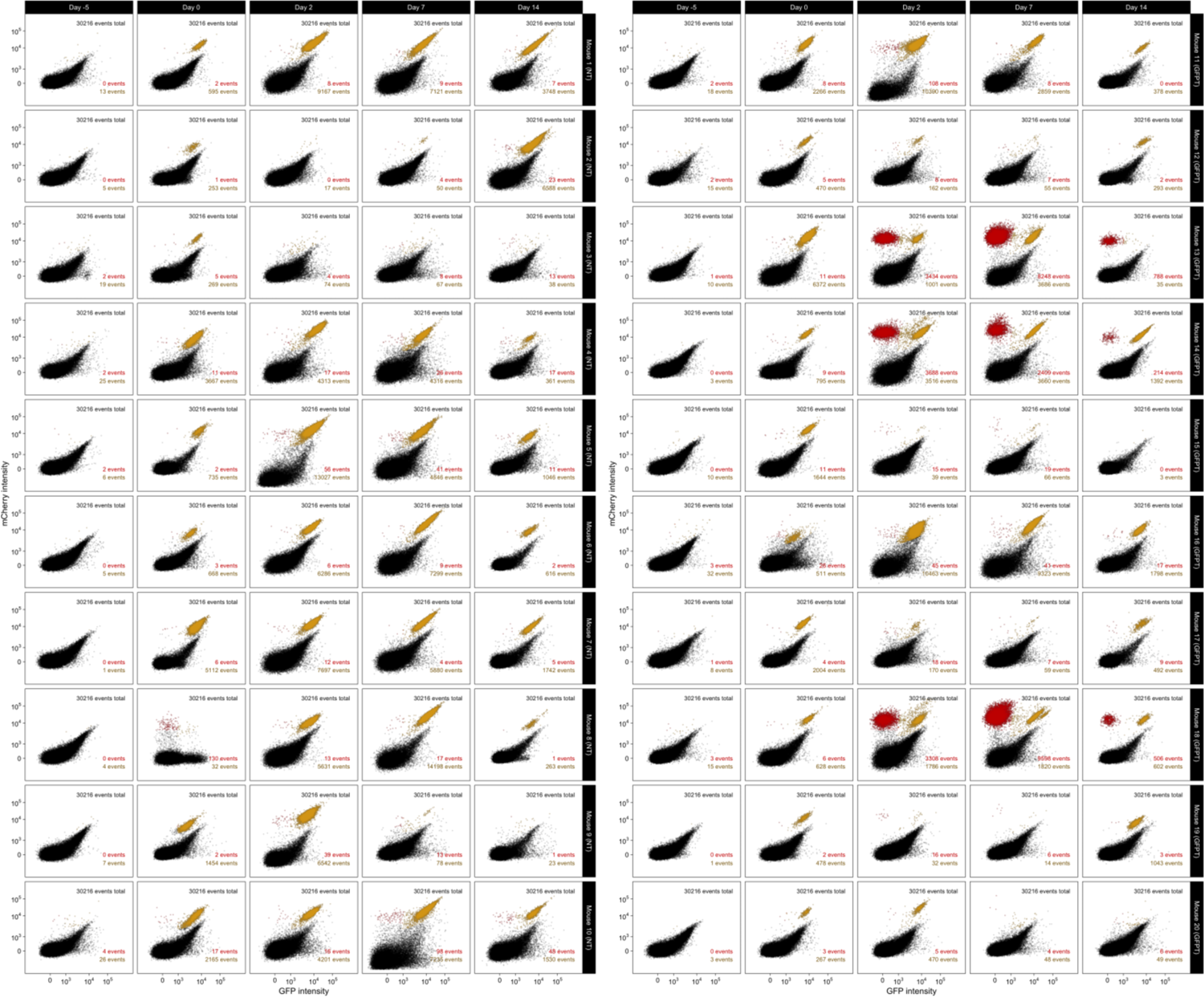
Flow cytometry plots of fecal samples for all mice at all timepoints during *in vivo* targeting of double-marked *E. coli*. Mice (n=10/group) were given either NT-M13 (left) or GFPT-M13 (right). Day −5, before colonization by *E. coli*; Day 0, after colonization by double-marked GFP+ mCherry+ *E. coli*; Day −2, post phage and carbenicillin treatment; Day 7, one week post-phage and carbenicillin; Day −14, one week after removing carbenicillin from drinking water. Based on visual inspection, the sample from Mouse 8 Day 0 was omitted from analyses.

**Supplementary Figure 16.**
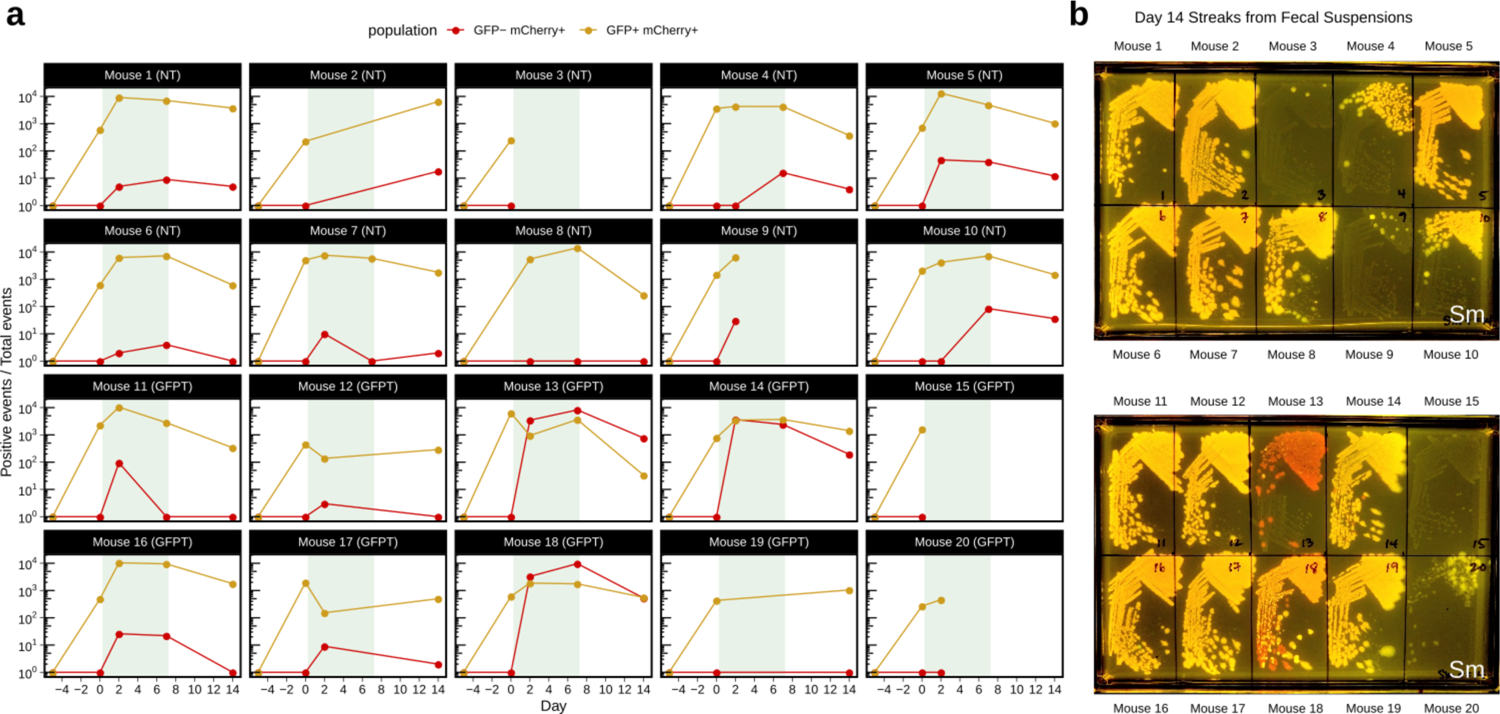
Fluorescence in fecal samples for individual mice colonized with double-marked GFP+ mCherry+ *E. coli* and dosed with NT-M13 or GFPT-M13. (a) For each mouse, the number of GFP+ mCherry+ events on Day −5 (before *E. coli* colonization) was used to subtract GFP+ mCherry+ background for all subsequent timepoints, and the number of GFP-mCherry+ events on Day 0 (before phage treatment) was used to subtract mCherry+ background from all subsequent timepoints. Shaded green area indicates duration of carbenicillin treatment. Timepoints were excluded when both GFP+ mCherry+ and GFP-mCherry+ events were below background thresholds. (b) Culture on LB with streptomycin from Day 14 fecal suspensions of mice treated with NT-M13 (M1 to M10, top) and GFPT-M13 (M11 to M20, bottom). Lack of fluorescent *E. coli* in fecal samples indicates eradication by carbenicillin where phage infection leading to colonization by Carb^R^ *E. coli* did not occur during the treatment phase.

**Supplementary Figure 17.**
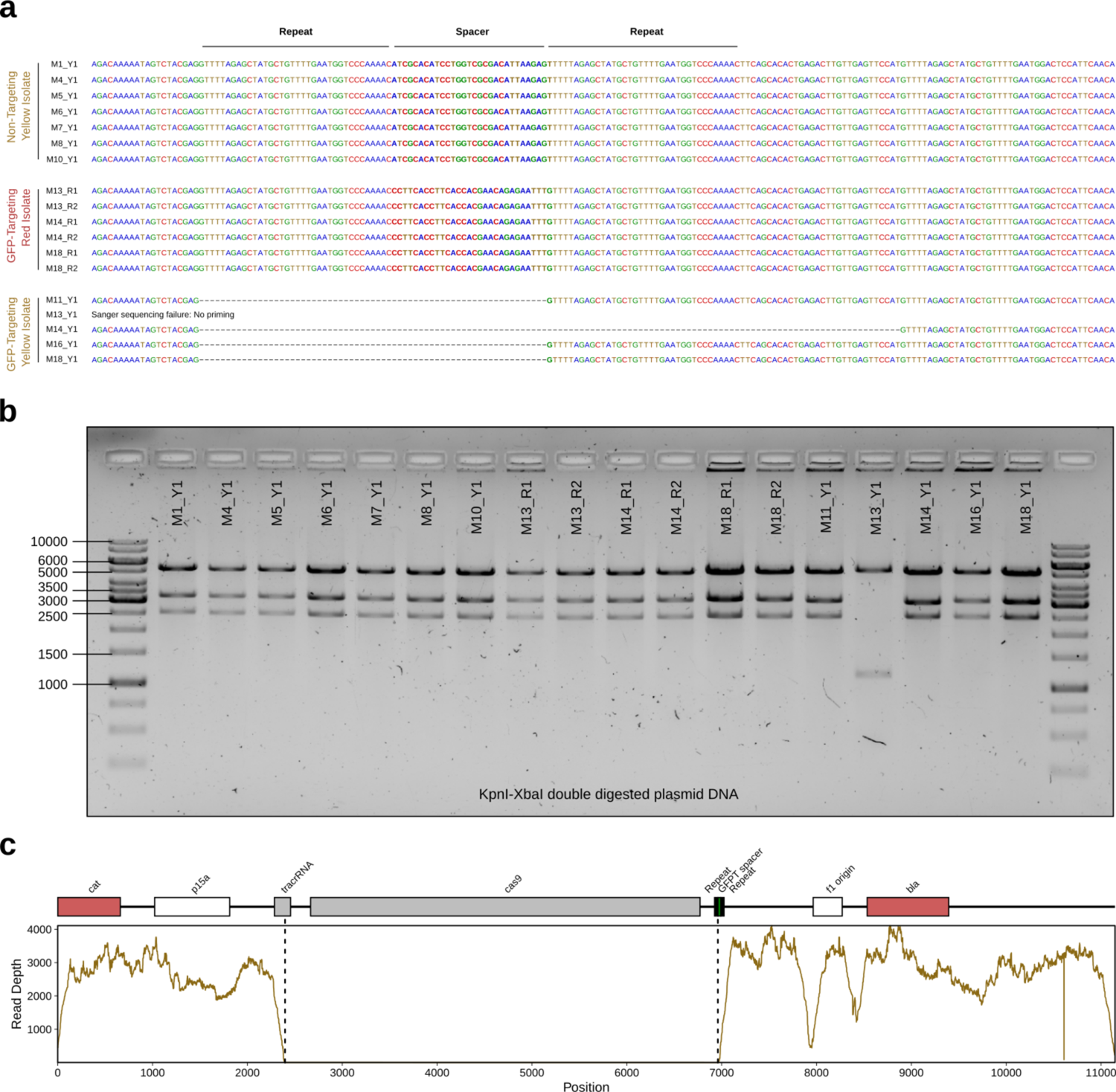
Mechanisms enabling double-positive *E. coli* to escape CRISPR-Cas9 targeting. (a) Sanger sequencing results confirm the expected spacer present in phagemid DNA extracted from fluorescent yellow isolates (Y1) colonizing NT mice (M1, M4, M5, M6, M7, M8, M10) and fluorescent red isolates (R1 and R2) colonizing GFPT mice (M13, M14, M18). In contrast, 4/5 fluorescent yellow isolates colonizing GFPT mice (M11, M13, M14, M16, M18) were confirmed to have lost the spacer. No Sanger sequence data was obtained for the last isolate (M13) with report for failing being “No Priming”, suggesting loss of a larger fragment from the phagemid. **(b)** Diagnostic digest of CRISPR-Cas9 phagemid DNA confirms a loss of phagemid DNA for the phagemid extracted from M13 Y1. Expected fragment sizes from KpnI-XbaI double digest: 5289, 3285, and 2573 bp. **(c)** Genome sequencing data for M13_Y1 also confirms loss of DNA from phagemid. Sequence coverage across the GFPT phagemid reveals lack of reads corresponding to the *cas9* gene, and parts of the CRISPR array and tracrRNA.

**Supplementary Figure 18.**
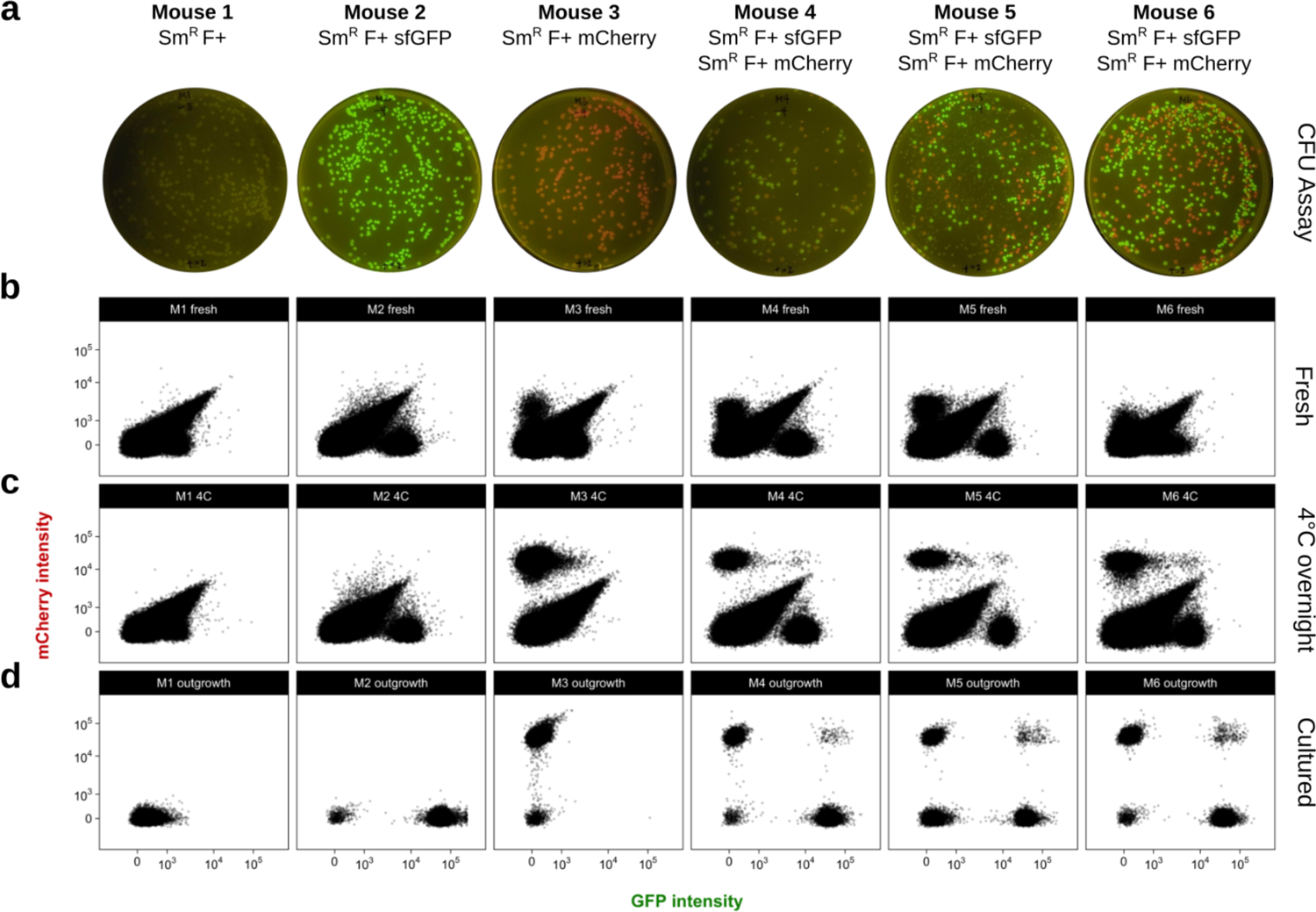
Quantification of fluorescent *E. coli* in mouse fecal pellets by flow cytometry improves with overnight incubation of fecal suspensions at 4°C. **(a)** Culture on LB streptomycin of fecal suspensions from streptomycin-treated mice colonized with non-fluorescent Sm^R^ *E. coli* W1655 F+ (Mouse 1), the GFP-marked (Mouse 2) or mCherry-marked derivative (Mouse 3), or both fluorescent strains (Mouse 4, 5, and 6). Flow cytometry was performed on fecal suspensions: **(b)** immediately after collecting, **(c)** overnight incubation at 4°C, or **(d)** after inoculation in media and overnight culture at 37°C.

## Supplementary Tables

**Supplementary Table 1.** Strains, plasmids, and phage.

**Supplementary Table 2.** Oligonucleotides.

